# Unveiling the sensory and interneuronal pathways of the neuroendocrine connectome in *Drosophila*

**DOI:** 10.1101/2020.10.22.350306

**Authors:** Sebastian Hückesfeld, Philipp Schlegel, Anton Miroschnikow, Andreas Schoofs, Ingo Zinke, André N Haubrich, Casey M Schneider-Mizell, James W Truman, Richard D Fetter, Albert Cardona, Michael J Pankratz

## Abstract

Neuroendocrine systems in animals maintain organismal homeostasis and regulate stress response. Although a great deal of work has been done on the neuropeptides and hormones that are released and act on target organs in the periphery, the synaptic inputs onto these neuroendocrine outputs in the brain are less well understood. Here, we use the transmission electron microscopy reconstruction of a whole central nervous system in the *Drosophila* larva to elucidate the sensory pathways and the interneurons that provide synaptic input to the neurosecretory cells projecting to the endocrine organs. Predicted by network modeling, we also identify a new carbon dioxide responsive network that acts on a specific set of neurosecretory cells and which include those expressing Corazonin (Crz) and Diuretic hormone 44 (DH44) neuropeptides. Our analysis reveals a neuronal network architecture for combinatorial action based on sensory and interneuronal pathways that converge onto distinct combinations of neuroendocrine outputs.

## Introduction

An organism is constantly subject to changing environmental challenges to homeostasis, and it counteracts these changes by adapting its physiology and behavior (Selye, 1956). In order to regulate homeostasis, animals must sense and integrate external and internal changes and generate outputs that comprise fundamental motivational drives such as feeding, fleeing, fighting and mating (Pribram, 1960). This output ultimately leads to motor activities through movement of muscles or through secretion of hormones that act on target tissues. The neuroendocrine system in any animal with a nervous system plays a vital role in controlling both forms of outputs. In its simpler form, e.g. in cnidarians, this takes place in a single sheet of epidermal cells that subsumes the functions of sensory, inter-, motor neurons and peptidergic cells (Grimmelikhuijzen et al., 1996; Martin, 1992). With more complex systems, the requirement for environmental sensing, integrating information and controlling motor outputs has given rise tospecialized neurons of the periphery and the central nervous system (Buijs and Van Eden, 2000; Hartenstein, 2006; Toni, 2004).

In mammals, different hormonal axes exist to keep essential physiological functions in balance, including the hypothalamic-pituitary-adrenal (HPA), the hypothalamic-pituitary-thyroid (HPT), the somatotropic and the two reproductive axes (Charmandari et al., 2005; Fliers et al., 2014; Grattan, 2015; Kaprara and Huhtaniemi, 2018). The various neuroendocrine axes also regulate each other. For example, the stress regulatory HPA axis relies on corticotropin releasing hormone (CRH) in the hypothalamus and has a negative influence on the reproductive regulatory axis (hypothalamic-pituitary-gonadal, HPG) by inhibiting gonadotropin releasing hormone (GnRH) (Kageyama, 2013; Rivier et al., 1986) such that when nutrients are scarce, the reproductive system is negatively affected until metabolic homeostasis is re-established (Tilbrook et al., 2002). The peptidergic basis for homeostatic regulation has also been characterized in invertebrates. These include, to name a few: stress (Johnson and White, 2009; Kubrak et al., 2016; Veenstra, 2009), metabolism and growth (Cannell et al., 2016; Dus et al., 2015; Gáliková et al., 2018; Geminard et al., 2006; Kahsai et al., 2010; Kim and Rulifson, 2004; McBrayer et al., 2007) and development (Hartenstein, 2006; Jindra et al., 2013; Truman, 2019; Truman et al., 1981; Wigglesworth, 1965). For comprehensive reviews, see (Nässel, 2018; Nässel and Winther, 2010; Nässel and Zandawala, 2019). Despite the extensive characterization of the neuroendocrine system in both vertebrates and invertebrates, very little is known regarding the sensory inputs to the neuroendocrine cells in the CNS. In general, a neuroendocrine system consists of neurosecretory cells in the brain that release peptides/hormones into the circulation to modulate effector organs. Via hormonal feedback loops, the neuroendocrine system is able to tune its regulatory function to set itself back into homeostasis. However, the synaptic pathways of sensory signals onto the neurosecretory cells in the brain remain largely elusive.

The *Drosophila* larva is a well-suited model to tackle the issue of the sensory pathways that act on the central neuroendocrine system. Parallels to the mammalian HPA system have been pointed out at physiological and anatomical levels. The pars intercerebralis (PI) and pars lateralis (PL) regions of the larval brain are considered to be analogous to the vertebrate hypothalamus. The three known endocrine glands (collectively known as the ring gland), the corpora cardiaca (CC), the corpus allatum (CA) and the prothoracic gland (PG) exert functions that are physiologically similar to the vertebrate pituitary gland (de Velasco et al., 2007; Hartenstein, 2006; Scharrer and Scharrer, 1944). These produce the critical metabolic, growth and maturation factors that are released directly into the circulation (adipokinetic hormone (AKH) from the CC; juvenile hormone (JH) from the CA; ecdysone from the PG). There are also analogies in basic functional and anatomical features that interconnect the hypothalamus and the brainstem in mammals, and the PI/PL region and the subesophageal zone (SEZ) in *Drosophila*. These also include the connections from the enteric nervous system to the CNS via the vagus nerve in mammals and the recurrent nerve in *Drosophila* (Schlegel et al., 2016; Schoofs et al., 2014b) (***Figure 1***).

**Figure 1.**
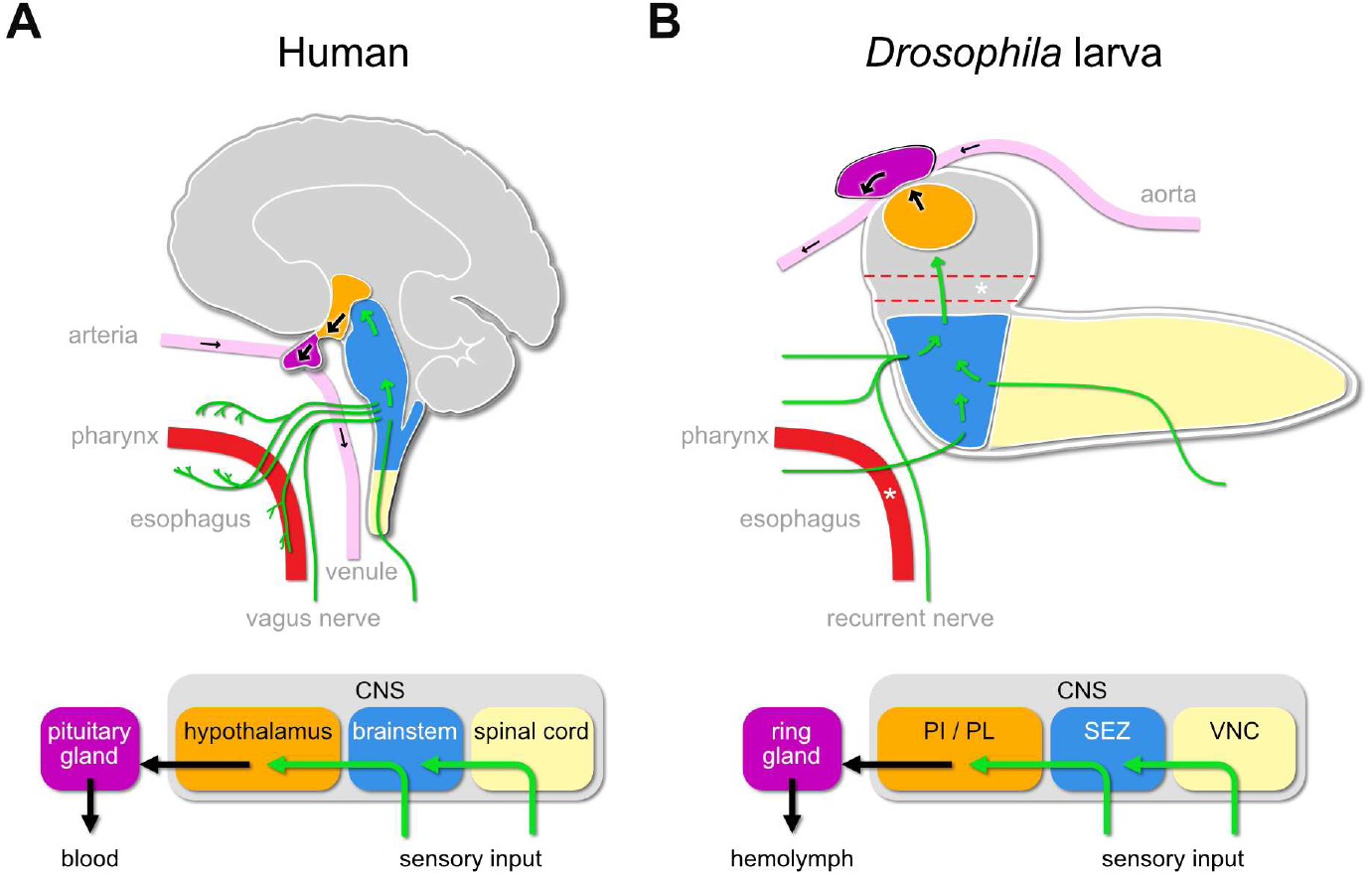
Sensory to endocrine pathways. (**A**) Schematic showing information flow from sensory input (green) to the endocrine system in the human brain compared to the *Drosophila* larval brain (**B**). PI: pars intercerebralis; PL: pars lateralis; SEZ: subesophageal zone; VNC: ventral nerve cord; CNS: central nervous system. Star denotes the foramen, where the esophagus (dotted red line) passes through the central nervous system. Green arrows denote flow of sensory information; black arrows denote release of hormones into the circulatory system.

Leveraging a synaptic resolution ssTEM volume of a whole L1 larval CNS (Eschbach and Zlatic, 2020; Miroschnikow et al., 2020; Ohyama et al., 2015; Schlegel et al., 2017; Thum and Gerber, 2019; Vogt, 2020), together with functional analysis of the hugin neuropeptide circuit, we have been characterizing the neuronal circuits that control specific aspects of feeding behaviour and the sensorimotor pathways of the pharyngeal nerves that drive food intake (Hückesfeld et al., 2016; Miroschnikow et al., 2018; Schlegel et al., 2016; Schoofs et al., 2014a). We now provide a comprehensive analysis of all neurosecretory cells that target the ring gland and the sensory neurons that form synaptic contacts with these cells, either directly or through interneurons. The neuronal network is organized in parallel interneuronal pathways that converge onto distinct combinations of neurosecretory cells in response to different sensory inputs. The circuit architecture allows variable and flexible action to maintain homeostasis and growth in response to broad multi-sensory and diverse metabolic signals. Using network modeling, we also identify novel CO_2_ responsive sensory pathways onto a specific set of neuroendocrine outputs.

## Results

### EM reconstruction of the neuroendocrine system

To elucidate the sensory inputs onto the neuroendocrine cells, we first reconstructed the ring gland and the interconnected portion of the aorta, and all neurons that project to these structures (***Figure 2A***). Reconstruction of a subset of the neurons in the PI was described earlier (Schlegel et al., 2016). All neurosecretory cell clusters found previously by light microscopy analysis (Siegmund and Korge, 2001) were identified and compared to expression patterns of known peptide-Gal4 driver lines. Since cell clusters that project to the ring gland (we collectively refer to them as Ring Gland Projection Neurons, RPNs) have varying names, we use here the peptide names that these neurons are mainly known for (***Figure 2 B, Figure 2 - figure supplement 1)***. CA-LP1 and CA-LP2 neurons were the only ones for which we could not unambiguously identify the neuropeptide identity, but found their expression in two independent Burs-Gal4 lines; also FMRFamide positive projections were found in the CA, which likely are derived from the CA-LP1 or CA-LP2 neurons (de Velasco et al., 2007). To analyse ion transport peptide (ITP) neurons (de Haro et al., 2010; Herrero et al., 2007; Kahsai et al., 2010), we generated LexA-knock-in lines (***Figure 2 - figure supplement 2***). A comprehensive overview for all RPN clusters analyzed in this study is provided as a supplementary table (***Figure 2 - source data 1***).

**Figure 2.**
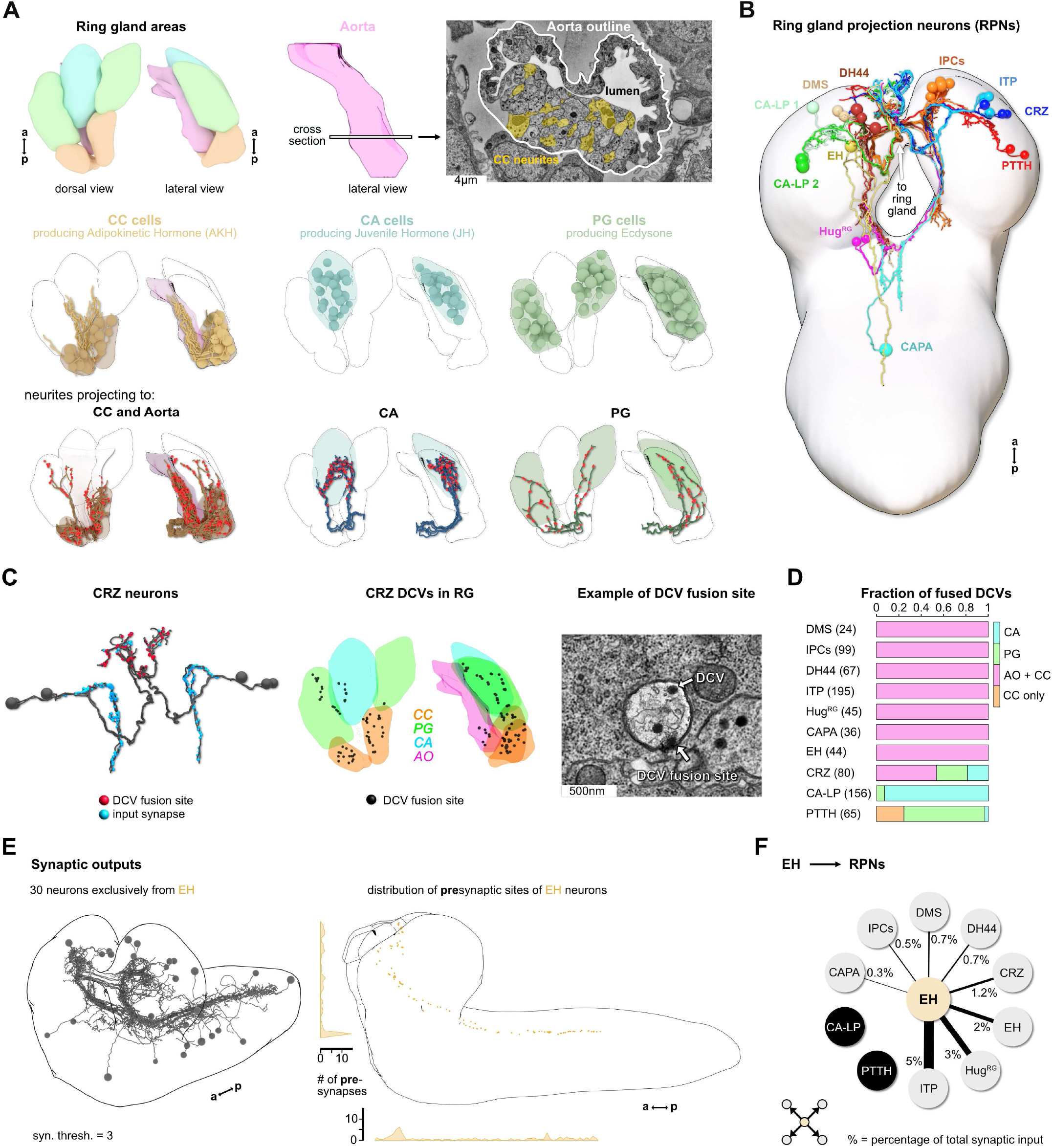
Reconstruction of the *Drosophila* larval ring gland and RPNs. (**A**) *upper panel*: 3D reconstructed ring gland areas in dorsal and lateral view (CC = orange, PG = green, CA = blue, Aorta = pink). Cross section of the aorta: Colored areas represent single neurites of different CC cells. *middle panel*: dorsal and lateral view of the ring gland showing the different cells in the distinct ring gland areas (CC, CA and PG). *lower panel*: Neurites innervating the RG areas were separated based on innervation of the CC and aorta, only CA or only PG. Fused dense core vesicles are marked as red dots. (**B**) Schematic of all neurons innervating the ring gland named by the main neuropeptide produced. For clarity, only one side is shown for each neuronal cluster. (**C**) *left*: Reconstructed CRZ neurons. Fused dense core vesicles were marked as non-polar output synapses at distal neurites in ring gland tissues (red dots). Blue dots represent chemical synaptic input sites. *middle*: Magnification of the reconstructed ring gland with spatial distribution of CRZ DCV fusion sites (black dots). *right*: example picture of a DCV fusion site in the EM volume (dense core vesicle has to be fused to the membrane). (**D**) Quantification of all DCV fusion sites found in the RG areas for each RPN group. Numbers in brackets are total numbers of marked DCVs. The X-axis represents a fraction of fused DCVs. (**E**) *left*: Synaptic outputs of all RPNs (threshold = 3 synapses) constitute in total 30 neurons, which are exclusively downstream of EH RPNs. *right*: Spatial distribution of presynaptic sites of EH. Neurons producing eclosion hormone are the only RPNs having presynaptic sites located along abdominal, thoracic segments and SEZ and Protocerebrum. (**F**) EH neurons synaptically target other RPNs. Percentage represents the fraction of input of distinct RPNs from EH neurons, e.g. ITP neurons receive 5% of its inputs from EH neurons. CC: corpora cardiaca; PG: prothoracic gland; CA: corpus allatum; RG: ring gland; IPCs: insulin producing cells; DMS: *Drosophila* myosuppressin; DH44: diuretic hormone 44; CRZ: corazonin; ITP: ion transport peptide; CA-LP: corpus allatum lateral protocerebrum; PTTH: prothoracicotropic hormone; HugRG: Hugin ring gland; CAPA: capability; EH: eclosion hormone; DCVs: dense core vesicles; RPN: ring gland projection neurons.

### Peptidergic and synaptic outputs

Peptidergic signaling is accomplished through release from dense core vesicles (DCVs). The specific peptidergic output region of all cells was identified by contacts of DCVs with the membrane with the apparent liberation of small dense particles, as exemplified for the Corazonin (Crz) neurons (***Figure 2 C***). The outputs of all ten peptidergic RPN groups are restricted mainly to the CC and aorta (AO). PTTH and CA-LP neurons project almost exclusively to the PG and CA, respectively (***Figure 2 D***). Neurons producing the stress related peptide Crz (Kubrak et al., 2016) showed the broadest output pattern, targeting all tissues (***Figure 2 C, D***). We also analysed the reliability of determining the output release site by quantifying DCV fusions sites. Using Crz and Crz receptor expressing cells as an example, we could confirm by trans-Tango system (Inagaki et al., 2012) that the CC cells are the main target of Corazonin neurons (***Figure 2 - figure supplement 3***). Thus, dense core vesicles found in the PG or CA might mean that other RPNs, like PTTH and CA-LP neurons, express the Crz receptor (for PTTH shown in (Imura et al., 2020). These data further lend support that DCV fusion sites represent a reliable measure for targets of RPNs. The anatomical data on peptidergic outputs were combined with existing single cell transcriptomic data on the larval brain (Brunet Avalos et al., 2019). Focusing on the expression of neuropeptides and their cognate receptors within the ring gland system, we confirmed for example that Crz neurons are targeting all other RPNs by releasing Corazonin as well as sNPF and Proctolin (***Figure 2 - figure supplement 4***). At the same time, the Crz receptor is expressed in the CC and to a lesser extent in the PG and CA, as well as in other RPNs. Based on the peptides and receptors expressed by the distinct RPN groups, the analysis uncovers complex interactions between neuroendocrine cells. At this point, it is unclear to what extent these peptide-receptor interactions occur between peptides released within the CNS or found in the hemolymph.

We next addressed the issue of the largely unknown synaptic connectivity of the neuroendocrine cells by reconstructing the synaptic up- and downstream partners of all RPNs (threshold of 3 synapses to each RPN). We identified 30 downstream partners which, unexpectedly, were exclusively targeted by the two eclosion hormone (EH) neurons, one on each side of the ventromedial protocerebrum (***Figure 2 E***). The functional significance of the EH synaptic outputs is as yet unknown. However, it has been shown that the neurohemal release sites could be removed and the axon stumps electrically stimulated; this evoked an ecdysis motor program through interaction of the EH neurons with response circuitry in the VNC (Hewes and Truman, 1991). Notably, these include all the other RPNs with the exception of CA-LP and PTTH neurons which regulate the activity of two major growth/maturation hormones, namely juvenile hormone (JH) and ecdysone (***Figure 2 F***).

### Synaptic inputs onto the neuroendocrine system

We identified 209 upstream partners of the RPNs, whose synaptic sites are distributed in the anterior thoracic and SEZ region and up along the protocerebrum in a sprinkler like fashion (***Figure 3 A***). Unlike the RPNs in the PI (IPCs, DMS, DH44), which have significant amounts of monosynaptic connections with sensory neurons (Miroschnikow et al., 2018; Schlegel et al., 2016), the RPNs of the PL (Crz, PTTH, CA-LP and ITP) have no direct sensory input. Similarly, EH, CAPA and Hug^RG^ RPNs have only small amounts of direct sensory contacts (***Figure 3 - figure supplement 1***). We therefore focused on the interneurons and their connection with the sensory system.

**Figure 3.**
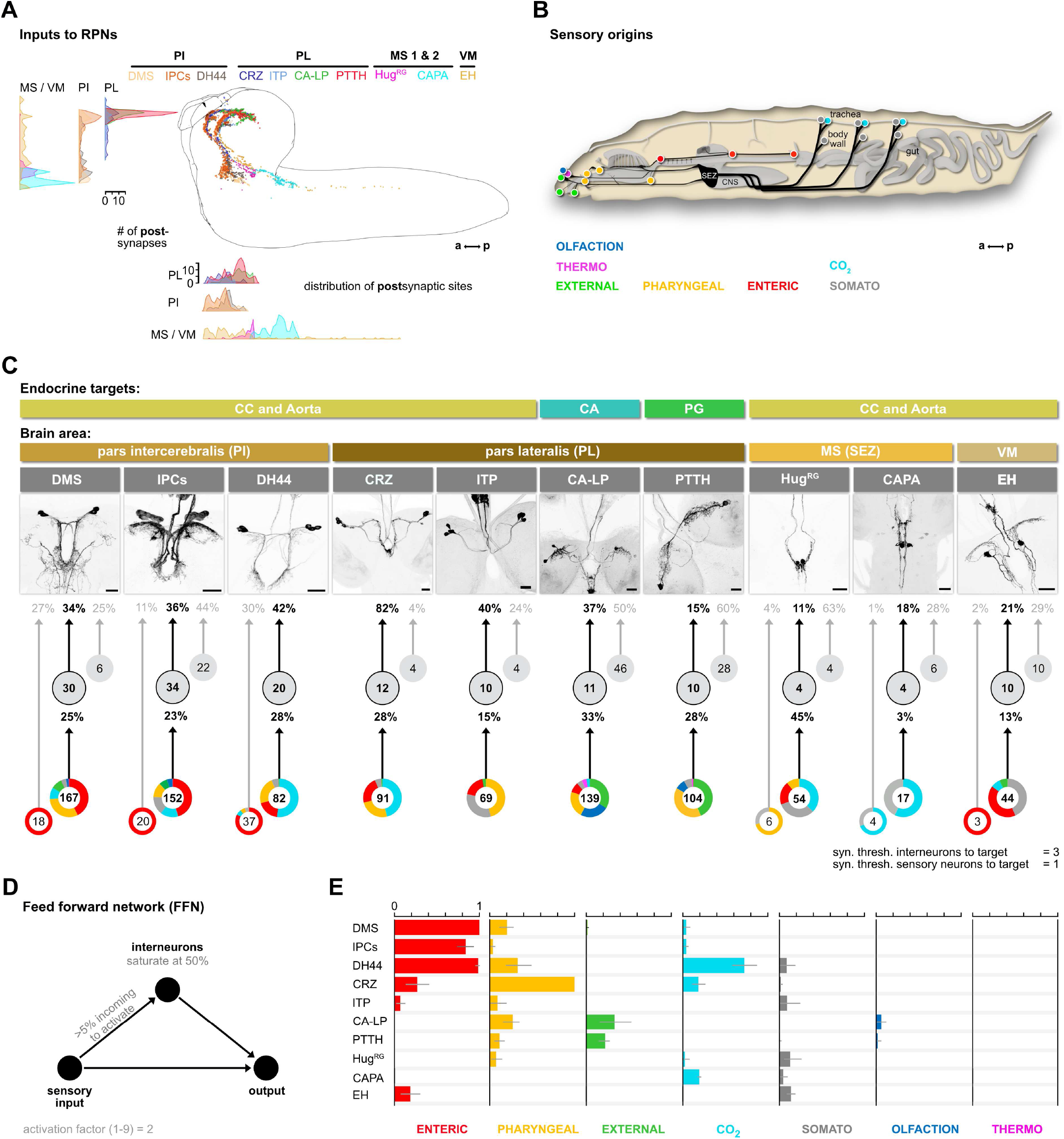
Inputs of RPNs and sensory origins. (**A**) Spatial distribution of postsynaptic sites of all RPNs (color coded). RPN postsynaptic sites are located along upper SEZ and in the protocerebrum in a sprinkler like fashion. (**B**) Schematic side view of a *Drosophila* larva. Colored dots represent the location of sensory organs, based on their sensory origin. (**C**) Synaptic connections to RPNs (grouped) *from top to bottom*: RPNs are grouped by their endocrine targets or their location of somata within the CNS (Brain area, colored bars). RPNs (displayed by expression pattern of the respective Gal4 or LexA lines) receive synaptic inputs (fraction of total synaptic inputs as percentage) from distinct sets of interneurons (numbers in circles represent the number of interneurons connected to RPNs), which in turn receive information from sensory neurons (fraction of total synaptic input as percentage). Colored pie charts represent the sensory profile of which interneuron groups of each RPN group integrate sensory neurons (numbers in white circles). Colors of pie charts correspond to the respective sensory origins shown in B. Note that the monosynaptic sensory neurons are also involved in polysynaptic pathways to the RPNs. (**D**) Scheme of the feed forward network (FFN). Sensory neurons are activated with an activation factor of 4 in the FFN. When more than 5% of presynaptic neurons are active, interneurons become activated up to an activity of 50%. (**E**) Summary of sensory driven modulation of RPN output groups by FFN. The X-axis for each panel shows the mean activity of RPNs listed on the Y-axis. Colors represent the different sensory origins used to activate the network through 1- and 2-hop synaptic connections.

We first divided the upstream interneurons into two groups: interneurons receiving direct sensory input and those that do not (threshold at 2 synapses); slightly more than half of all upstream neurons integrate sensory information, n=110 (***Figure 3 - figure supplement 1***). Based on previous publications, we know the peripheral origin (e.g. enteric, pharyngeal, olfactory) of most sensory neurons (Berck et al., 2016; Miroschnikow et al., 2018). Here, we additionally characterize a subset of tracheal dendritic neurons (TD neurons) (Schlegel et al., 2016) as being responsive to CO_2_ (***Figure 3 - figure supplement 2***). To determine which sensory signals are integrated by RPNs via these interneurons, we grouped their sensory inputs based on their peripheral origin (***Figure 3 B***). The resulting map provides a comprehensive overview of the sensory to endocrine pathways in the larval neuroendocrine system (***Figure 3 C***). All of the RPNs receive input from a distinct combination of interneurons, which in turn receive input from a distinct combination of sensory neurons. In one extreme (e.g. IPCs), 152 sensory neurons from six different sensory regions (greatest from enteric) target 34 interneurons. At the other extreme (e.g. CAPA), 17 sensory neurons from two sensory regions target just 4 interneurons. The synaptic load of RPNs from interneurons that receive sensory inputs vary greatly. The largest is for Crz, where 82% (fraction of input synapses) of the total input is from interneurons with direct sensory connections.

### Modelling the impact of activating sensory neurons on the neuroendocrine system

To assess the potential impact of sensory inputs on the neuroendocrine system, we employed a network diffusion model based on direct monosynaptic and 2-hop polysynaptic connections using feed forward connectivity (***Figure 3 D***). The model is deliberately kept simple as we lack detailed knowledge on the physiology (e.g. neurotransmitter) of the neurons involved. Such networks have been recently used successfully in the mouse to model sensory impact on activity in higher brain centers of the thalamus (Shadi et al., 2020). Our model predicts the impact of specific sensory origins onto each RPN group (***Figure 3 E***, for parameterization and connection types in the model see ***Figure 3 - figure supplement 3.1 and 3.2***; adjacency matrix for all neurons used in this study in ***Figure 3 - figure supplement 4***). As a first experimental analysis based on the predictions, we chose the CO_2_ path because the defined sensory organ, i.e. tracheal dendritic (TD) neurons, and distinct modality (CO_2_) made it more tractable.

### A novel CO_2_ to endocrine pathway

The model predicts a strong impact of TD neurons on DH44, Crz, DMS and CAPA RPNs (***Figure 4 A***). To validate this, we performed imaging experiments using the ratiometric calcium integrator CaMPARI-2 to measure changes in activity of the RPNs upon CO_2_ exposure. Indeed, the in-vivo experiments confirmed the predictions: DH44, Crz and DMS RPNs were strongly activated by CO_2_ (***Figure 4 A*** - right panel, experimental details in ***Figure 4 - figure supplement 1***). CAPA neurons did not differ significantly from control groups but tended to show a lower activity upon CO_2_ stimulation. Since the network diffusion model does not take the sign of a connection into account, it is conceivable that CAPA neurons are inhibited by CO_2_. The analysis of connectivity based on the EM volume enabled us to identify a new circuit in which CO_2_ is detected by TD neurons, integrated by a core set of four thoracic interneurons (somata located in T1-T3 segments) which then in turn strongly connect to DH44 and Crz neurons (***Figure 4 B***). Each of the thoracic interneurons have slightly different connectivity profiles in terms of their up and downstream partners (***Figure 4 C***). Thus, while all four are interconnected to CO_2_ sensory neurons and target DH44 or Crz neurons, the strength of the connections differs as well as their connections to other sensory and RPN neurons. Please see ***Figure 4 - figure supplement 2.1 and 2.2*** for identity (ID number and connectivity) of all interneurons.

**Figure 4.**
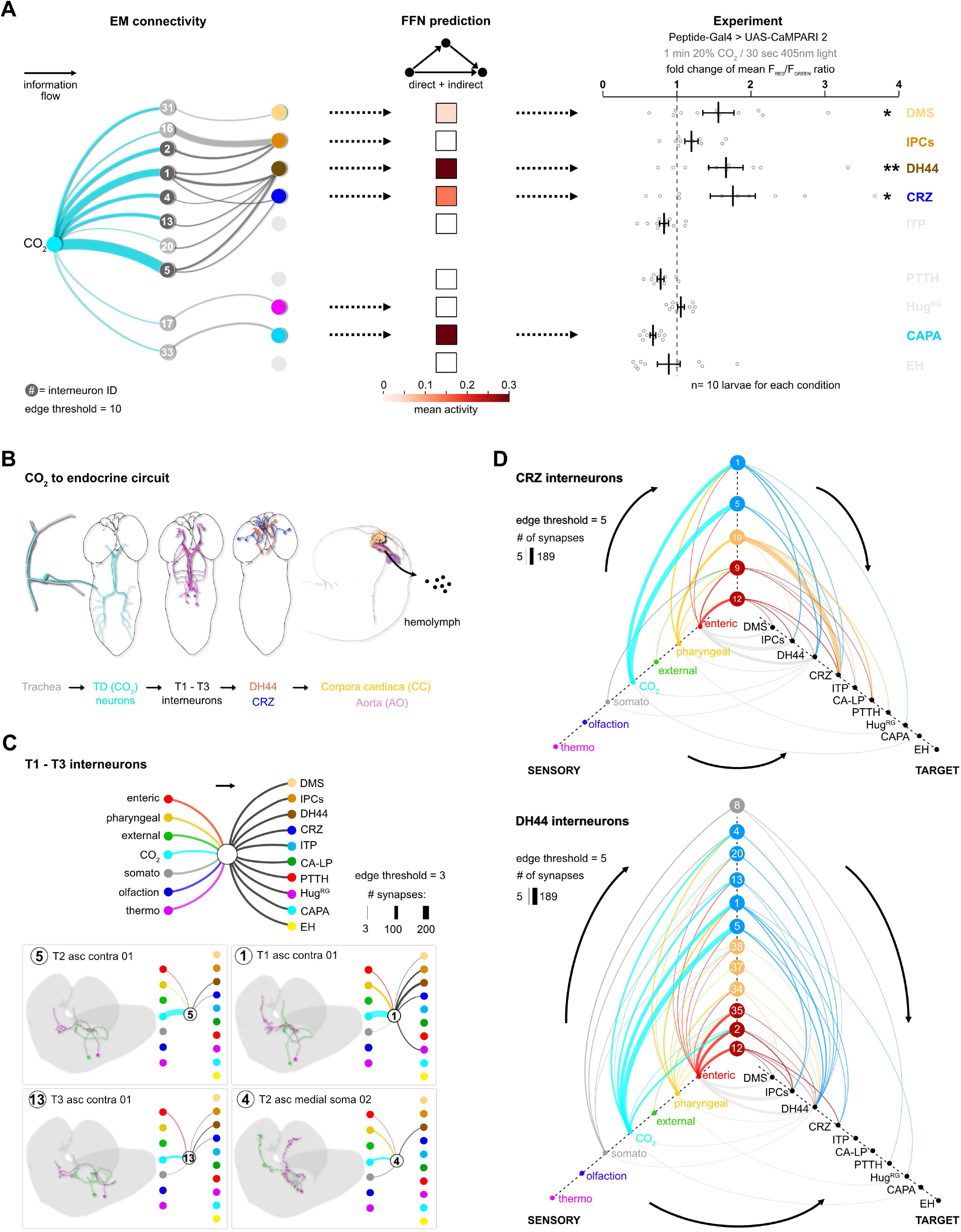
CO_2_ pathway from TD neurons to RPNs. (**A**) Comparison of underlying connectivity of TD CO_2_ neurons via interneurons to the RPNs, with the predicted outcome of mean activity (with an activation factor of 4) of RPNs, and the outcome of CaMPARI-2 CO_2_ experiments. Network diffusion model reliably shows modulation of the RPNs. (**B**) Using the combination of connectivity, prediction and functional imaging experiments, a new sensory to endocrine neural circuit can be derived. TD CO_2_ neurons sense CO_2_ at the trachea and communicate predominantly via a core set of thoracic interneurons to DH44 and CRZ expressing cells, which show release sites in the CC and aorta. (**C**) Connectivity of the single thoracic interneurons (hemilateral pairs) to presynaptic sensory origins and to the distinct postsynaptic RPN groups. Thoracic interneurons receive additionally other sensory modalities apart from CO_2_ neurons, and target different combinations of RPNs. (**D**) CRZ interneurons: Hive plot showing the polysynaptic pathways from all sensory origins to all RPN target groups, using the interneurons (synaptic threshold = 3) which target CRZ neurons. Main sensory origins are enteric, pharyngeal and CO_2_. DH44 interneurons: CO_2_ neurons represent the most dominant polysynaptic path from sensory origins to DH44 neurons. Note that monosynaptic connections from sensory neurons to RPNs are shown in grey.

We then took the two main output RPNs of the CO_2_ circuit (Crz and DH44) and asked what other interneurons were upstream of these, and to which sensory neurons these interneurons were connected (***Figure 4 D***). For Crz, the strongest are in fact not the thoracic interneurons from the CO_2_ pathway: one hemilateral pair of interneurons (#10, Munin 2) accounts for over 50% of total synaptic input to the Crz neurons. These interneurons receive sensory information exclusively from pharyngeal sensory neurons (***Figure 4 D***, top hive plot). There are two other strongly connected interneurons (#9, Munin 1; #12, SiB) and they receive most of their inputs from the enteric region. Furthermore, all the interneurons are also part of pathways that target several RPNs. For example, interneuron #10 targets all neurons of the PL, whereas interneuron #12 targets all neurons of the PI. For DH44, the strongest upstream partners are the same thoracic interneurons that respond to CO_2_ (***Figure 4 D***, bottom hive plot). In sum, this illustrates the distinct sensory-to-neuroendocrine connectivity profiles (which sensory origins onto which set of RPNs) of the different interneurons. For summarized connectivity schemes of the RPNs, see ***Figure 3 C***.

### Interneurons that direct sensory information to distinct sets of neuroendocrine outputs

We next extended the connectivity hub analysis to the other interneurons of the neuroendocrine system (***Figure 5 A***). Therefore, we calculated the fraction of sensory inputs to given interneurons and multiplied it with the fraction of inputs of the RPN. This analysis revealed interneurons that play a major role in the sensory pathways to the neuroendocrine system. Selected notable interneurons are illustrated in ***Figure 5 B***. For example, both Hugin PC (#11) and SiB (#12) interneurons have their strongest inputs from the enteric sensory neurons; however, whereas Hugin PC interneurons strongly target just the IPCs (edge threshold of minimum 5 synapses), SiB interneurons target DMS, IPCs and Crz (***Figure 5 B***).

**Figure 5.**
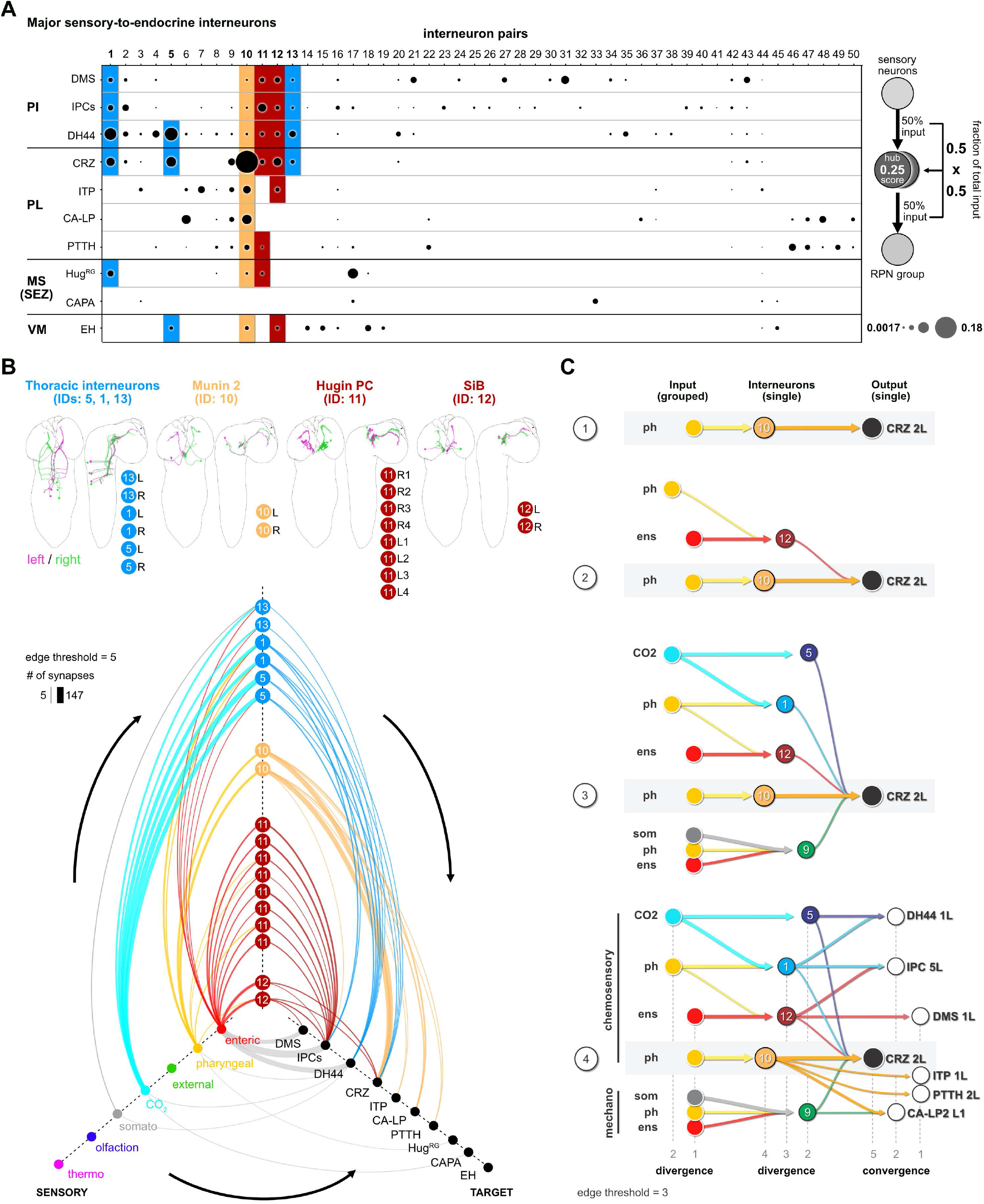
Circuit architecture of sensory to endocrine pathways. (**A**) Dot plot showing the importance of interneurons acting as sensory to endocrine hub. Dot size was calculated using the fraction of total input an interneuron receives from sensory neurons multiplied by the fraction of total input this interneuron gives to an RPN output group. Colored backgrounds of dots are highlighted in B. (**B**) Selected interneurons (highlighted in A) connecting the sensory system with RPNs. Thoracic interneurons receive sensory information from TD CO_2_ neurons and target IPCs, DH44 and CRZ (hive plot, strongest connection= 147 synapses). Munin 2 interneurons connect CRZ, ITP, PTTH and CA-LP RPNs with pharyngeal sensory neurons. Hugin PC neurons connect the IPCs with enteric sensory neurons. SiB neurons also receive information from enteric origins but target DMS, IPCs and CRZ. Edge threshold for hive plot = 5 synapses. (**C**) Circuit architecture common for all RPNs (CRZ single cell example). **1**: the strongest polysynaptic path based on hub analysis from pharyngeal sensory origin to CRZ output neuron via interneuron 10. **2**: Second interneuron (12) integrating enteric information and different pharyngeal information, converging onto CRZ output neuron. **3**: all interneurons of one CRZ output neuron integrating multiple sensory origins and converging onto one single output. **4**: Concept of divergence and convergence in the neuroendocrine connectome. Sensory neurons diverge / converge onto distinct sets of interneurons. Interneurons diverge in varying synaptic strength onto distinct RPN output neurons. Numbers at bottom show degree of convergence and divergence (e.g. interneuron 10 diverges to CRZ, ITP, PTTH and CA-LP. All interneurons converge to CRZ. Synaptic threshold = 3 for all connections).

There are also intriguing unique groups, e.g. the interneurons (#s 46-50) which are highly specialized for CA-LP and PTTH (***Figure 5 A***; ***Figure 5 - figure supplement 1***); these receive strong sensory inputs from the olfactory system (for a comprehensive connectivity map see ***Figure 5 - figure supplement 2***). In adult *Drosophila* it was shown that the release of juvenile hormone from the CA potentiates sensitivity of a pheromone sensing olfactory receptor OR47b (Lin et al., 2016) to maximize courtship success of male flies. In larvae, we found several previously appetitive and aversive assigned ORs (Kreher et al., 2008) being connected via multi-glomerular projection neurons to the CA-LP and PTTH neurons. This might be relevant for larvae where ecdysone or juvenile hormone would be secreted in response to olfactory cues, although the function of such a pathway is not known. For a summary of interneurons connecting to each RPNs, see ***Figure 5 - figure supplement 3***.

Finally, we illustrate the key features of the neuronal circuit architecture that underlie the neuroendocrine system, which can be constructed using Crz as an exemplary RPN (single output cell) (***Figure 5 C***). We start with the strongest connection from interneuron Munin 2 (#10), which receives input from a group of pharyngeal sensory neurons (***Figure 5 C, panel 1***). A second interneuron SiB (#12) receives input from a group of enteric sensory neurons (***Figure 5 C, panel 2***); this interneuron also receives inputs from a different class of pharyngeal sensory neurons. More interneurons are added to build a series of parallel paths (diverging sensory signals) that all converge on a common RPN (***Figure 5 C, panel 3***). This set of interneurons concurrently target different RPNs (***Figure 5 C, panel 4;*** see also ***figure legend*** for details). Thus, the parallel paths that converge on a single RPN (e.g., Crz) additionally target multiple RPNs. For single cell networks of all RPNs, see ***Figure 5 - figure supplement 4***.

## Discussion

### The neuroendocrine connectome of *Drosophila* larvae

Organisms differ in their adaptive capacity to deal with external and internal changes, but the essential goal remains the same: ensuring homeostasis in a changing environment. Evolution of neuroendocrine systems led to the separation of sensory systems, neuroendocrine cells and specialized glands (Hartenstein, 2006). We show in this paper how the central neuroendocrine system is synaptically organized. A general feature of the ring gland projection neurons (RPNs) is the absence of synaptic outputs within the CNS. The exception is the eclosion hormone (EH) producing neurons, which have synaptic outputs in the protocerebrum, SEZ and ventral nerve cord. This unique feature of EH neurons might be due to their function in coordinating movements during larval cuticle shedding (Baker et al., 1999; Krüger et al., 2015). Another feature is that the RPNs of the PL are connected with the sensory organs exclusively via polysynaptic paths, which is in contrast to the numerous monosynaptic connections found for RPNs of the PI (Miroschnikow et al., 2018; Schlegel et al., 2016). It is also noteworthy that peptides known for their roles in metabolic and stress regulation in general receive large amounts of their inputs from interneurons with direct contacts to the sensory system, i.e. these paths are short, with only a single hop between the interneurons and sensory neurons. This might be due to the need for rapid action, compared to those (e.g. PTTH and CA-LP neurons) involved in gradual, long term and irreversible events such as larval growth and maturation.

### Novel CO_2_ responsive sensory to endocrine pathways: from connectomic based modeling to in vivo testing

Numerous previously unknown synaptic pathways from the sensory organs to the RPNs were revealed from our connectomic analysis, including a new set of sensory neurons, namely the tracheal dendritic CO_2_ neurons (TD CO_2_) that respond to CO_2_. This might be due to the stress associated with high levels of CO_2_, which is observed in humans as well (Permentier et al., 2017). These sensory neurons target, via thoracic interneurons, RPNs that express two peptides known to play a dominant role in metabolic stress regulation in *Drosophila*: Diuretic hormone 44 (DH44) and Corazonin (Crz) (Cannell et al., 2016; Dus et al., 2015; Kubrak et al., 2016). From a neuronal network perspective, it was possible to predict this modulation with a feed forward network. Both peptide groups display homology to mammalian neuroendocrine axes known to regulate stress (HPA axis) and reproductive behaviour (GnRH axis). DH44 is a homolog of vertebrate corticotropin releasing hormone (CRH), which is released in the hypothalamus in response to external and internal stressors like hypoxia or hypoglycaemia (Flanagan et al., 2003). A role for DH44 in glucose and amino acid sensing has been reported (Dus et al., 2015; Yang et al., 2018), but its role in CO_2_ sensing was not previously known. CO_2_ activation of Corazonin, a homolog to GnRH, adds to the repertoire of stress sensors ascribed to these neurons that include their roles in glucose and fructose sensing (Dus et al., 2015; Kubrak et al., 2016; Miyamoto and Amrein, 2014; Oh et al., 2019; Veenstra, 2009). The connectome analysis further indicates that Crz and DH44 neurons have the strongest synaptic connections with the sensory system (i.e., greatest number of paths that are connected monosynaptically or via single interneurons), suggesting a critical role of these neurosecretory cells in rapid sensory integration.

### Combinatorial parallel pathways enable variability and flexibility in the central neuroendocrine system

Sensory pathways are often studied based on a single type of sensory organ or modality, in most cases for technical reasons. In a natural environment, it is unlikely that an animal will encounter a situation where it needs to react to only a single sensory input and secrete a single type of hormone. For the fly larvae, two broad types of actions have to be taken into account: immediate action to an acute stress (e.g. due to toxic smoke, predator wasp or starvation), and a slower action that enables tissue and organismal growth (e.g. accumulation of biosynthetic resources for cell growth and progression onto the next moulting or puparium stage). Even an acute response takes place within the existing physiological state of the organism. For the endocrine organs, this requires the secretion of different combinations, and most likely different concentrations, of hormones and neuropeptides into the circulation or target tissues.

At the core of the neuroendocrine network is a parallel set of interneurons that target distinct combinations of neuroendocrine outputs (e.g. the RPNs, each expressing certain neuropeptides). Each of the interneurons in turn receive sensory inputs from distinct sets of sensory neurons (e.g. CO_2_ sensitive in trachea or different type and modality within the pharyngeal region). This can be also seen in the pathways from olfactory sensory neurons to CA-LP and PTTH endocrine targets. Multi-glomerular projection neurons integrate olfactory as well as gustatory information, and as one proceeds deeper into the neuronal circuitry, interneurons that have originally been classified as interneurons without sensory input can be connected by additional hops to sensory neurons (such as through mushroom body and lateral horn in the protocerebrum). These then converge together with the multi-glomerular projection neurons onto the common set of interneurons that target the CALP and PTTH output neurons. The different converging paths can be seen to represent distinct types of sensory information, including a stored form from the mushroom body (Eichler et al., 2017; Eschbach and Zlatic, 2020; Miroschnikow et al., 2020) where a positive or negative valence has been attached to an existing sensory cue. Additionally, there are a significant number of synaptic connections among the interneurons. Such architecture would enable variability and flexibility in the combination and concentrations of neuropeptides that become released in response to the flood of multi-sensory inputs that act on all parts of the neuroendocrine network. Subsequently cross-regulatory interactions at the receptor level would then determine the final neuropeptide/hormone composition that is released within the CNS or into the circulation. Our work provides a neuronal architectural blueprint of how this is constructed at the synaptic level for the neuroendocrine system in the brain and may also be of general relevance in understanding other complex neuroendocrine systems.

As a concluding remark, the neuroendocrine connectome of the *Drosophila* larva presented here (i.e. the “ring gland connectome”) represents the first complete synaptic map of sensory to endocrine pathways in a neuroendocrine system of this complexity, and adds another level of insight on the known humoral functions of the released neuropeptides and hormones. Together with the large amount of knowledge on the function of neurosecretory cells targeting the CC, CA, PG and aorta over the past years (Nässel and Zandawala, 2020), the current analysis increases our understanding of how the neuroendocrine system receives information about external and internal sensory cues. A future challenge in this context is the identification of specific sensory neurons of different origin and modality to define the valence of sensory integration, and the function of the interneurons that enable different pathways to the endocrine organs.

## Acknowledgments

We thank the whole Pankratz lab for fruitful discussions on earlier versions of this manuscript. We thank Anna Pepanian, Christina Georgopoulou und Stephan Nottelmann for help with cloning of the ITP-T2A-LexA line. We thank Dick Nässel, Michael B. O’Connor, Jae Park, Yi Rao, Jan Veenstra, Christian Wegener and Benjamin White for sharing fly lines and resources. We also thank all “tracers” for their contribution to the EM reconstruction.

## Materials and Methods

### Flies

All larvae used for experiments and stainings were 96 +/- four hours (after egg laying) of age and were grown on standard cornmeal medium under a twelve hour light/dark cycle if not otherwise stated. Following Gal4 driver and UAS effector lines were used:

*Ilp2-Gal4* (IPC neurons, BL#37516), *Ms-Gal4* (DMS neurons, (Park et al., 2008)), *Dh44-Gal4* (DH44 neurons, BL#51987), *Crz-Gal4* (CRZ neurons, BL#51977), *CrzR-Gal4*^*T11A*^ (Sha et al., 2014), *Ptth-Gal4* (PTTH neurons, (McBrayer et al., 2007), *Burs-Gal4* (BL#51980), *Burs-Gal4* (BL#40972, this line shows expression in CA-LP neurons of the PL, data not shown), *Eh-Gal4* (EH neurons, BL#6301), *17B03-Gal4* (Hug^RG^ neurons, (Jenett et al., 2012), *714-Gal4* (CAPA neurons, (Gohl et al., 2011)), *ITP-T2A:Gal4* (ITP neurons, used in CaMPARI experiments, unspecific expression in CNS glia observed BL#84702), *ITP-T2A:LexA* (ITP neurons, for generation see below, used in stainings - clean expression of ITP in the CNS), *260-Gal4* (TD CO_2_ neurons, BL#62743), *UAS-mRFP* (BL#27398), *UAS-CaMPARI-1* (BL#58761), *UAS-CaMPARI-2* (BL#78316), *trans-Tango* (BL#77124), *lexAop2-myrGFP* (BL#32209).

**Table:**
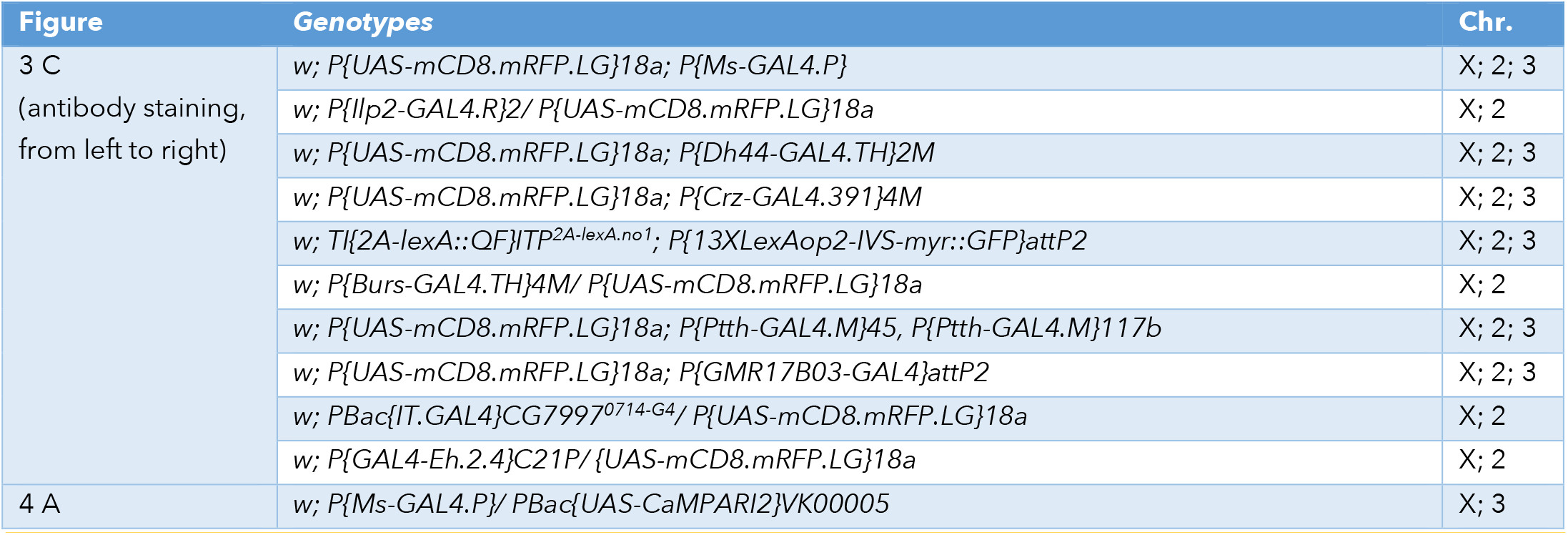
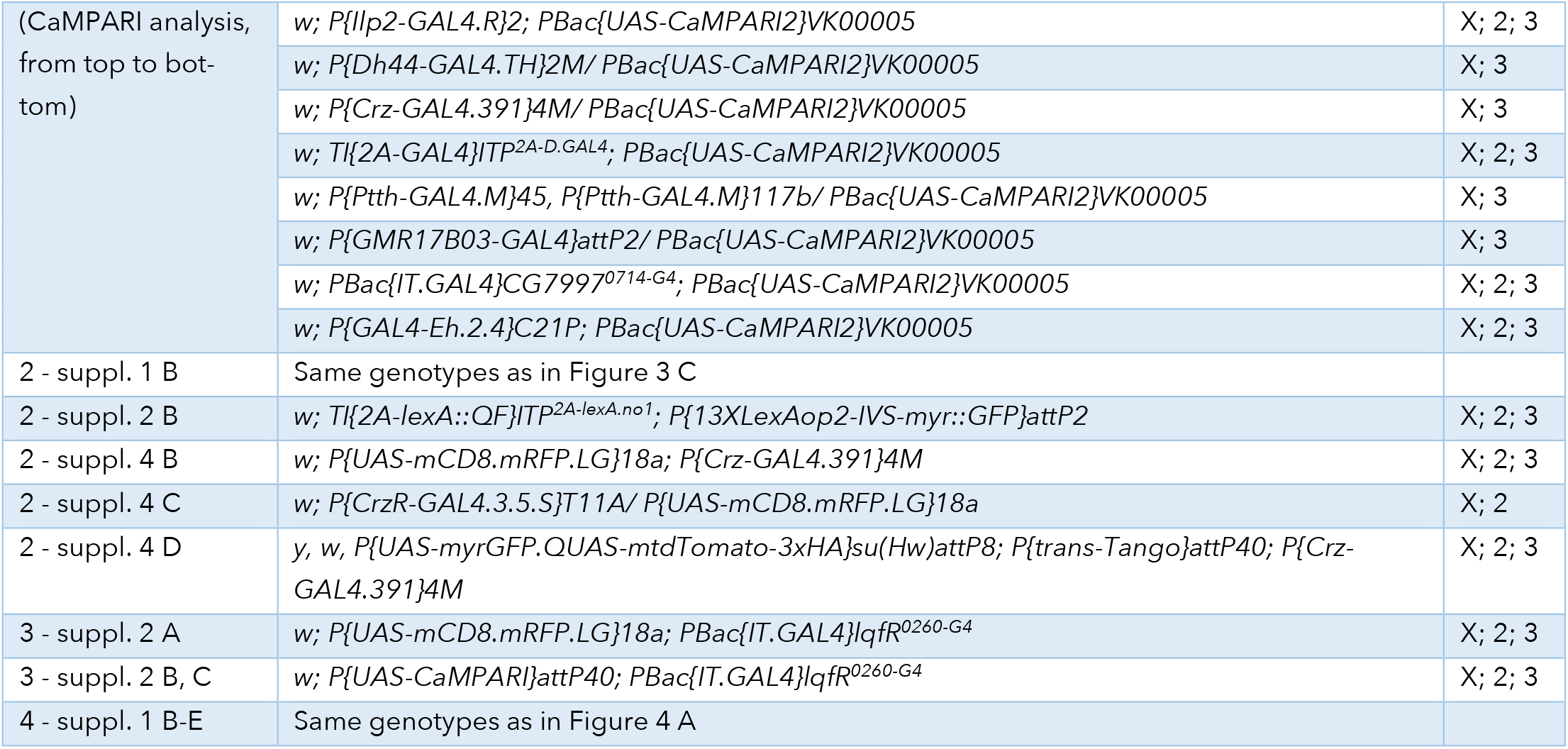
Genotypes of experimental flies.

### Generation of *ITP-T2A-LexA* transgenic fly lines

First we generated *T2A-LexA:QF* knock-in constructs that can be targeted to genomic loci by homology-directed repair using the CRISPR/Cas system. Therefore, *splice acceptor-T2A-LexA:QF* fragments for all three intron phases were amplified by PCR (Q5 polymerase, New England Biolabs) from *pBS-KS-attB2-SA(0/1/2)-T2A-LexA::QFAD-Hsp70* plasmids (Addgene #62947, #62948 and #62949) (Diao et al., 2015) with primers CGTACTCCACCTCACCCATC and ctcgagAAGCTTCTGAATAAGCCCTCGT. PCR products were sub-cloned into *pCRII-TOPO* vector (invitrogen) to create plasmids *TOPO-T2A-LexA:QF(0), TOPO-T2A-LexA:QF(1)* and *TOPO-T2A-LexA:QF(2)*. Next *splice acceptor-T2A-Gal4* cassette from *pT GEM(0)* (Addgene #62891) (Diao et al., 2015) was removed by *Xba*I/*Sal*I digest and replaced with *Xba*I/*Xho*I fragments from *TOPO-T2A-LexA:QF(0), TOPO-T2A-LexA:QF(1)* and *TOPO-T2A-LexA:QF(2)* harboring *splice acceptor T2A-LexA:QF* cassettes (T-LEM, T2A-LexA expression module) for all three intron phases. All restriction enzymes used and T4 DNA ligase are from New England Biolabs. We named these *T2A-LexA:QF* knock-in plasmids *pT-LEM(0), pT-LEM(1)* and *pT LEM(2)*.

Two CRISPR target sites (no1 and no2) in the intron downstream of the first coding exon shared by all five predicted transcripts of the *Ion transport peptide* gene (*ITP*) to insert T-LEM were chosen using *flyCRISPR Optimal Target Finder* (*Gratz et al*., *2014*). By ligating annealed oligonucleotides two guide RNA expression constructs were inserted into *Bbs*I-linearized *pCFD3* vector (Port et al., 2014). Sequences of oligonucleotides were

(no1) gtcgGTGTTCCTTACAGCGTTCA aaacTGAACGCTGTAAGGAACAC

(no2) gtcgAAAATGATCGCGGGACCTT aaacAAGGTCCCGCGATCATTTT.

Next, 5prime and 3prime homology arms (5’HA, 3’HA) for both targeted sites were introduced into *pT-LEM(2)*. Therefore, target site flanking sequences of approximately 1kb size were amplified by PCR (Q5 polymerase, New England Biolabs) from genomic DNA of *nos-Cas9*^*[attP2]*^ fly line used for embryo injection. Primer sequences see in Table below. PCR products were sub cloned into *pCRII-TOPO* vector (invitrogen). Then 5’HAs were ligated as *Sph*I/*Not*I fragments from TOPO plasmids into *Sph*I/*Not*I-linearized *pT-LEM(2)* vector resulting in *pT-LEM(2)-5’HA-no1* and *pT-LEM(2)-5’HA-no2*. Finally 3’HA no1 was inserted as *Asc*I/*Kpn*I fragment from TOPO plasmid into *Asc*I/*Kpn*I-digested *pT-LEM(2)-5’HA-no1* and 3’HA no2 as *Kpn*I/*Spe*I fragment into *Kpn*I/*Spe*I-cut *pT-LEM(2)-5’HA-no2*, resulting in *pT-LEM(2)-ITP-no1* and *pT-LEM(2)-ITP-no2*, respectively. Plasmid microinjections to generate *ITP*^*T2A-LexA-no1*^ and *ITP*^*T2A-LexA-no2*^ lines were performed by BestGene Inc. By using Cre-loxP system the 3xP3-DsRed cassette was removed from *ITPT2A-LexA-no1* and *ITP*^*T2A-LexA-no2*^.

**Table:**
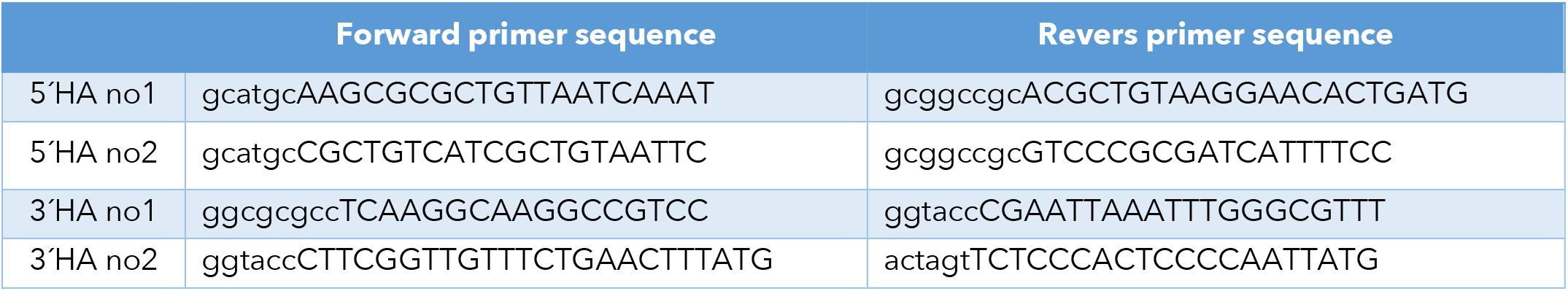
Primer sequences to generate homology arms.

### EM reconstruction

Neuron reconstruction was done on an ssTEM (serial section transmission electron microscope) volume of a six hour old first instar larva (Ohyama et al., 2015). We identified the RPNs by reconstruction of all axons originating in the CNS and targeting the ring gland through the NCC nerve. The mNSCs including neurons producing Insulin-like peptides, DMS and DH44 have been previously reconstructed and described (Miroschnikow et al., 2018; Ohyama et al., 2015; Schlegel et al., 2016). We reconstructed all neurons to completion (tracing 100% and at least 95% reviewed). Downstream targets were not synaptically connected to RPNs (except for EH downstream partners, being reconstructed with a synaptic threshold of 3). Therefore, membrane fused dense core vesicles were marked as connectors without direction. Dense core vesicles within the CNS were not marked due to technical issues with the common synapse annotation system. No synaptic connections were observed in the larval ring gland. The ring gland was reconstructed with all cells and tissue areas were assigned based on tissue boundaries, color (CA area was slightly darker, CC cells showed dendritic arborizations into the CC) and cell soma position. All synaptic up- and downstream partners of the RPNs were reconstructed to completion with a synaptic threshold to each of the RPNs of three synapses.

For sensory neurons included here, we made use of earlier published data (Berck et al., 2016; Miroschnikow et al., 2018; Ohyama et al., 2015; Schlegel et al., 2016). A subset of twelve TD neurons were already described (Schlegel et al., 2016). We reconstructed for this study all 26 TD neurons.

### Sensory neuron pie charts

Pie charts in Fig. 3 and following: Pie charts of sensory profiles were calculated using the percentage of total synaptic input of interneurons and RPNs (in case of monosynaptic connections) as fraction (thereby ignoring other inputs to show distribution of sensory origins). Percentages then give the percentage of total sensory synaptic input to interneurons or RPNS.

### Hubscore

Calculation of Hub Score in Fig. 5A: Fraction of total synaptic input from all sensory neurons to defined interneurons (see IDs) was multiplied by the total fraction of input of the RPN group from this interneuron. E.g. Interneuron #10 (Munin2) receives 32.33% (fraction: 0.3233) of their total synaptic input from sensory neurons. In turn, Corazonin neurons receive 56.52% (fraction: 0.5652) of their total synaptic inputs from interneuron #10 (Munin2). Multiplying the fractions of this path (sensory via interneuron to CRZ) leads to a hub score of: 0.3233 × 0.5652 = 0.18272916 (hub score).

### Immunohistochemistry

Dissected larval brains were fixed for one hour in paraformaldehyde (4 %) in 1x phosphate-buffered saline (PBS), rinsed three times (20 min) with 1% PBS-T (1% Triton X-100 in 1x PBS) and blocked in 1% PBS-T containing 5% normal goat serum (ThermoFisher) for one hour. Primary antibody was added to the solution (for concentrations, see below). Brains rotated overnight at 4°C. On the second day larval brains were washed three times (20 min) with 1% PBS-T and subsequently secondary antibody was applied. Brains rotated overnight at 4°C. After three times washing with 1% PBS-T brains were dehydrated through an ethanol-xylene series and mounted in DPX Mountant (Sigma-Aldrich). Imaging was carried out using a Zeiss LSM 780 confocal microscope with 25x or 63x objective (oil). For antibody stainings of *peptide>mRFP*, the primary antibody was anti-RFP (1:500, mouse, abcam, ab65856). Secondary antibody was anti-Mouse Alexa Fluor 568 (1:500, goat, Invitrogen, A-11031). For *ITP>myr-GFP* stainings primary antibody was anti-GFP (1:500, chicken, abcam, ab13970) and secondary antibody was anti-Chicken Alexa Fluor 488 (1:500, goat, Invitrogen, A-11039). For Crz staining primary antibody was anti-Crz (1:500, rabbit, gift from C. Wegener), secondary antibody was anti-Rabbit Alexa Fluor 568 (1:500, goat, Invitrogen, A-11011). For Trans-Tango stainings primary antibodies were anti-GFP (1:500, chicken, abcam, ab13970) and anti-RFP (1:500, mouse, abcam, ab65856). Secondary antibodies were anti-Chicken Alexa Fluor 488 (1:500, goat, Invitrogen, A-11039) and anti-Mouse Alexa Fluor 568 (1:500, goat, Invitrogen, A-11031), respectively. For staining of nuclei of the ring gland, diamidino-phenylindole (Roth) was used 1:1000.

### Functional Imaging with CaMPARI

For experiments with TD-neuron line *260-Gal4* we used *UAS-CaMPARI1* (Fosque et al., 2015). A larva was placed inside the petri dish and fixed with duct tape for 60 seconds. 405 nm UV light (M405L2_UV, Thorlabs) was placed twelve cm above the larva and illuminated with a LED controller (LEDD1B, Thorlabs at max intensity) for 15 seconds. Afterwards the larval brain was dissected and put onto a poly-L-lysine coated coverslip and covered with 1x PBS for imaging at low Ca^2+^ conditions. Caudal dendrites of TD neurons which project to the SEZ were imaged. For defined concentrations of CO_2_ stimulation, we used a CO_2_ incubator (CB 53, Binder) at CO_2_ concentrations of 0, 10 and 20% CO_2_ at 24-27°C. Stimulation protocol was the same as described before.

For experiments with different *peptide-Gal4* lines we used *UAS-CaMPARI2* with improved baseline fluorescence and improved integration dynamics (Moeyaert et al., 2018). In our hands photoconversion ratios were lower in general but more defined when neurons were not active, lowering the number of false positive photoconversion (own observations). We used the CO2 incubator to set CO_2_ concentration to 20% and compared neuronal photoconversion with 0% CO_2_ concentration in the incubator. Larvae were placed on duct tape in the middle of a five cm petri dish for 60 seconds and afterwards illuminated for 30 seconds with 405 nm at max intensity. Following steps were the same as described before.

### Statistics

For CaMPARI experiments green to red ratios of single cells of *peptide-Gal4* lines were analyzed with a custom-made script for FIJI (ImageJ) and the mean was calculated per animal (each cell was analyzed and a mean build). Animal means were then analyzed and plotted with Prism 6 software using the Mann-Whitney-Rank-Sum test, p<0.05*, p<0.01**, p<0.001***, p<0.0001****.

### Feed forward network diffusion model

The feedforward network (FFN) was implemented in Python as a simple artificial neural network without backpropagation. Synaptic weights were normalized by the total number of post-synapses such that they represented fractions of inputs for a given neuron. Neurons were implemented as rectified linear units using a ReLu activation function that starts responding at 5% and reach saturation at 50% of their synaptic inputs being active:

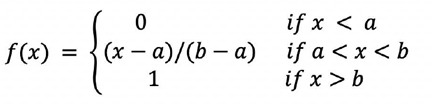

With x being the sum activity of all inputs weighted by their synaptic weights, constants a and b controlling the response onset and saturation, respectively. a and b were chosen such that neurons start responding at 5% and reach saturation at 50% of their synaptic inputs being active: a=0.05, b=0.5. These values were chosen to maximize the response range of the network. The code for the FFN and the generation of the figures can be found at https://github.com/Pankratz-Lab/FFN_Hueckesfeld-et-al.-2020.

### Analysis of single cell transcriptomic data from Brunet Avalos et al. 2019

In order to analyze peptide receptor interaction between RPN groups we sought out to use the data generated in the lab of Simon Sprecher describing the single cell transcriptomic atlas of the *Drosophila* larval brain (Brunet Avalos et al., 2019). Advantage of this dataset was the exclusive analysis of SEZ and brain lobes, which helped in finding the RPN specific peptidergic cell groups. We used R analysis similar to the described workflow in (Brunet Avalos et al., 2019), based on Seurat v3 workflow (Butler et al., 2018; Stuart et al., 2019). In brief, we used seurat processing pipeline from Satija lab (https://satijalab.org/seurat/) to process the integrated datasets of fed and starved conditions (GEO accession number GSE134722 (Brunet Avalos et al., 2019). This combined dataset consists of 9346 cells and 14064 analyzed features. In order to cluster the RPNs into the specific groups, following parameters were used: dataset: fed and starved integrated and log normalized | scale = 10,000 | 2000 variable genes | Seurat v3 processing: Cells with unique features: 200 - 4500 | genes expressed in at least 1 cell | 31 PCs were used to assess cell clusters | resolution was 1 | cluster 12 was identified as peptidergic cells | Peptidergic cells were separated with following parameters (expression profiles):

**IPCs**: Ilp2 >=3 & Ilp5 & Ilp3 (26 cells)

**DMS:** Ms >=6 (9 cells)

**DH44:** Dh44 >=2.8 (12 cells)

**CRZ:** Crz>=1 &sNPF >=1 (13 cells)

**ITP:** ITP >=1 & Lk>=0.8 (17 cells)

**PTTH:** Ptth >=2 (9 cells)

**CA-LP:** FMRFa >3.5 (14 cells)

**Hug**^**RG**^: Hug > 4 & Mip >1 (7 cells)

**CAPA:** Capa >=2 (6 cells)

**EH:** Eh >=4 (4 cells)

For CA-LP neurons FMRFamide was used based on the description in de Velasco et al. 2007. Hugin-RG cells were separated based on Coexpression of Mip neuropeptide (unpublished observation, staining with Mip-Gal4 line and Hugin-antibody).

### Graphic representation and visualization

Neurons were rendered in Blender 3D (ver2.79b) using the CATMAID to Blender interface described by (Schlegel et al., 2016) (https://github.com/elifesciences-publications/Catmaid-toBlender) and edited in Affinity Designer (Serif) for MAC. Staining images were processed with FIJI (ImageJ) and CaMPARI images were analyzed using a custom made FIJI script to be subsequently edited in Affinity Designer. Hive Plots were generated by using the CATMAID software for spatial distribution of nodes and subsequently made in Gephi 0.92 with rescaled edge weights (e.g. 1-200 synapses were rescaled for line thickness 1-20). Edges with less than five synapses were ignored in Gephi. To visualize peptide receptor connectivity, we used Circos tableviewer (http://mkweb.bcgsc.ca/tableviewer/).

## Notes

### Competing Interest Statement

The authors have declared no competing interest.

