## Supplementary file for "Unveiling the sensory and interneuronal pathways of the neuroendocrine connectome in *Drosophila*"

### Figure supplements

**Hückesfeld et al**

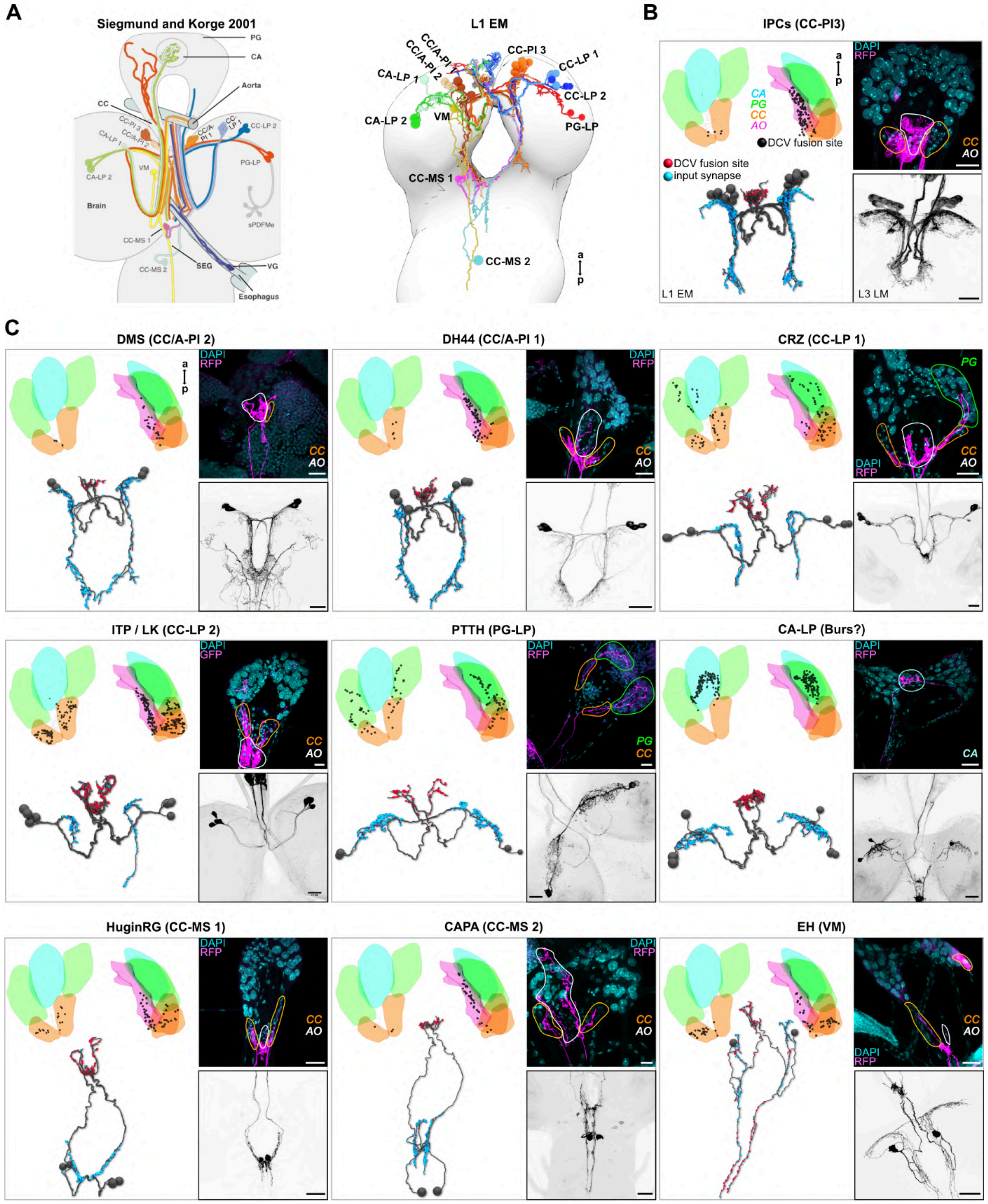


**Figure 2 – figure supplement 1. Comparison between L3 light microscopy and larval L1 ssTEM data of RPNs.** (**A**) All RPNs could be re-identified by comparing the neurons innervating the ring gland previously published by (Siegmund and Korge, 2001) to all RPNs found in the L1 ssTEM volume (right panel). (**B,C**) DCV fusion sites were analyzed (black dots in dorsal and lateral view of the ring gland) in the L1 ssTEM dataset (upper left part of each panel). Lower left part of each panel: blue dots represent chemical post-synapses in the brain and red dots represent DCV fusion sites in the RG. Note that pre-synapses for EH neurons are also found in the CNS (red dots). RPNs identified in the ssTEM volume were compared to respective expression patterns of Gal4/lexA lines in fluorescence stainings (UAS-mRFP/lexAop-myr-GFP, right side of each panel). Upper right of each panel displays Gal4 driven expression patterns of RFP/myr-GFP in the ring gland co-stained with DAPI to visualize RG areas.Note that CA-LP neurons were visualized using a Bursicon-Gal4 line. This expression in the CA has not been reported yet for Bursicon.

A

B

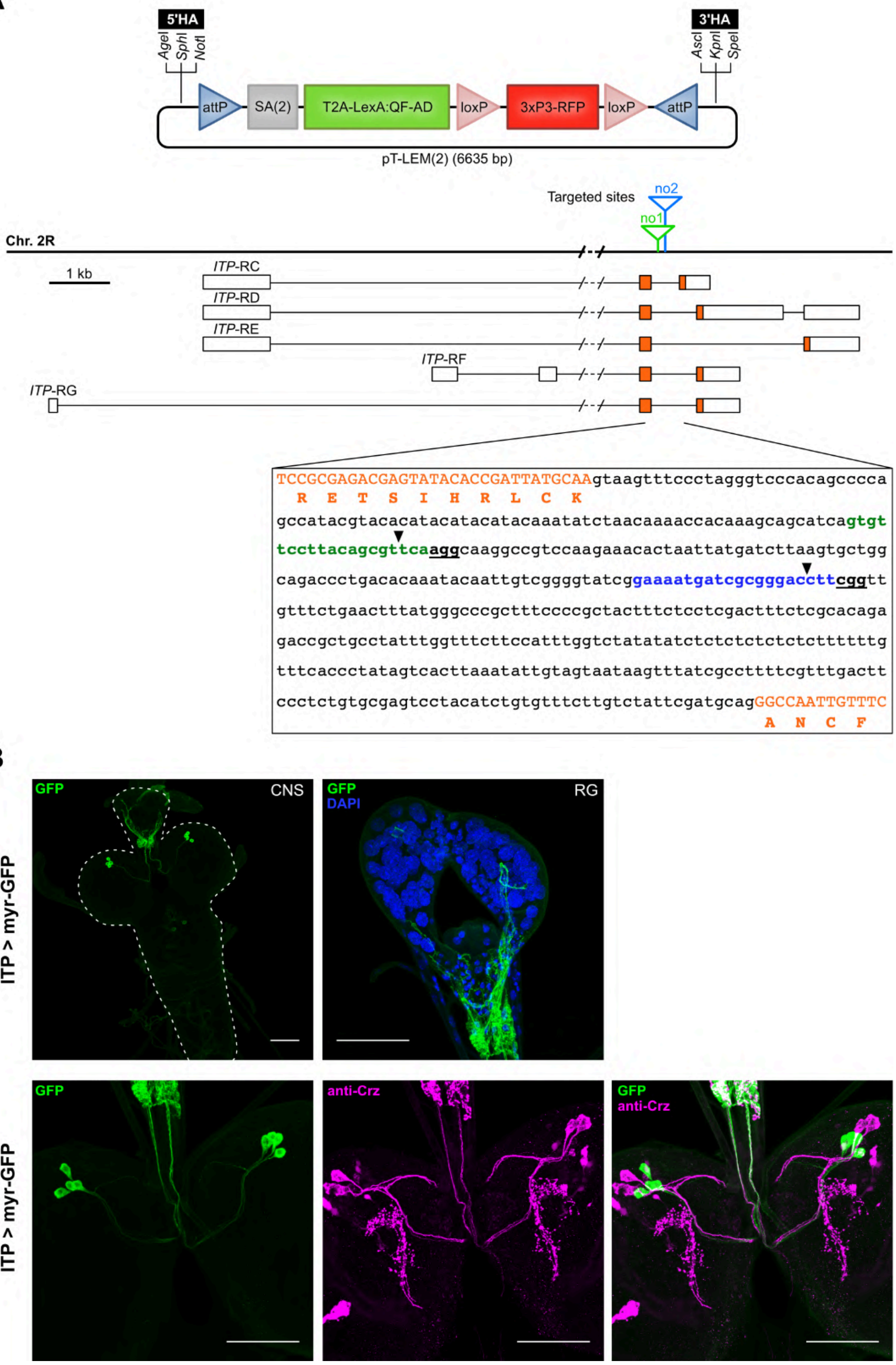


**Figure 2 - figure supplement 2. CRISPR/Cas dependent integration of T-LEM cassettes into *ITP* intron.** (**A**) Schematic of the T2A-LexA expression module (T-LEM) vector *pT-LEM(2)*, which contains red fluorescent protein (RFP) under the control of synthetic eye promoter 3xP3. Multiple cloning sites flanking T-LEM permit insertion of appropriate homology arms to target *ITP* intron. Splice acceptor sequence and phase 2 linker (SA(2)) in *pT-LEM(2)* allows integration of T-LEM as new exon into phase 2 introns. Genomic locus on the right arm of 2nd chromosome (2R) with all five predicted splice variant transcripts of *ITP* gene depicting two intronic sites (no1, no2) targeted for T-LEM insertion. Boxes represent exons; orange areas show protein-coding regions. Sequence of *ITP* intron displaying protospacers no1 (green) and no2 (blue). Protospacer adjacent motifs (PAM) are bold and underlined. Coding exons are capitalized and orange. One letter code amino acid sequence is shown in orange and positioned under the first nucleotide of the encoding triplet. Black arrowheads mark target sites for T-LEM insertion. (**B**) Fluorescence staining of ITP-T2A-LexA line driving lexAop-myr-GFP. upper panel: expression in whole CNS and ring gland. lower panel: expression of ITP-T2A-LexA line (green) and colocalization with anti-CRZ antibody reveals no overlap of both RPN groups.

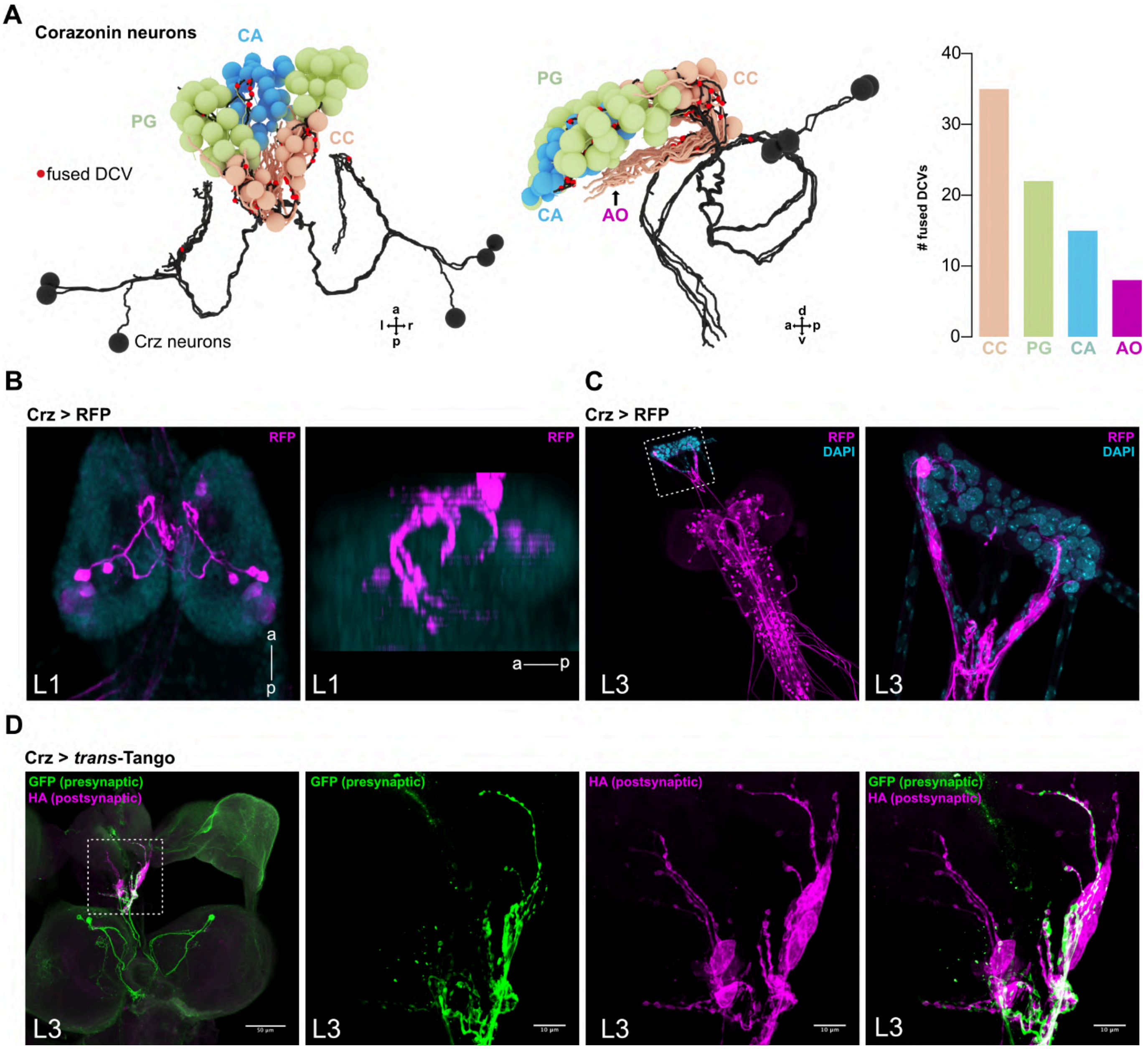


**Figure 2 - figure supplement 3. Fused DCV sites as proxy for real release sites in the ring gland: example CRZ**. (**A**) Reconstructed CRZ neurons in dorsal and lateral view with projections to RG cells (red dots are DCV fusion sites). Right bar plot shows quantification of the number of fused DCVs in CC, PG, CA and AO. Most output release sites were found in the CC. (**B**) Analysis of the Crz-Gal4 line in first instar (L1) larval CNS shows only activity in CRZ neurons projecting to RG. View angles are the same as in (A). (**C**) Analysis of Crz receptor Gal4 line (CrzR-Gal4) in third instar (L3) larval CNS (left) and in RG (right) shows CrzR activity in CC cells. (**D**) Staining of post-'peptidergic' cells of Crz expressing neurons using the *trans*-Tango system (for details see Materials and Methods section; note: no output synapses for Crz in L1 ssTEM volume. Whether synapses might be present in the L3 stage is not known). Only CC cells were marked as being postsynaptic to CRZ RPNs.

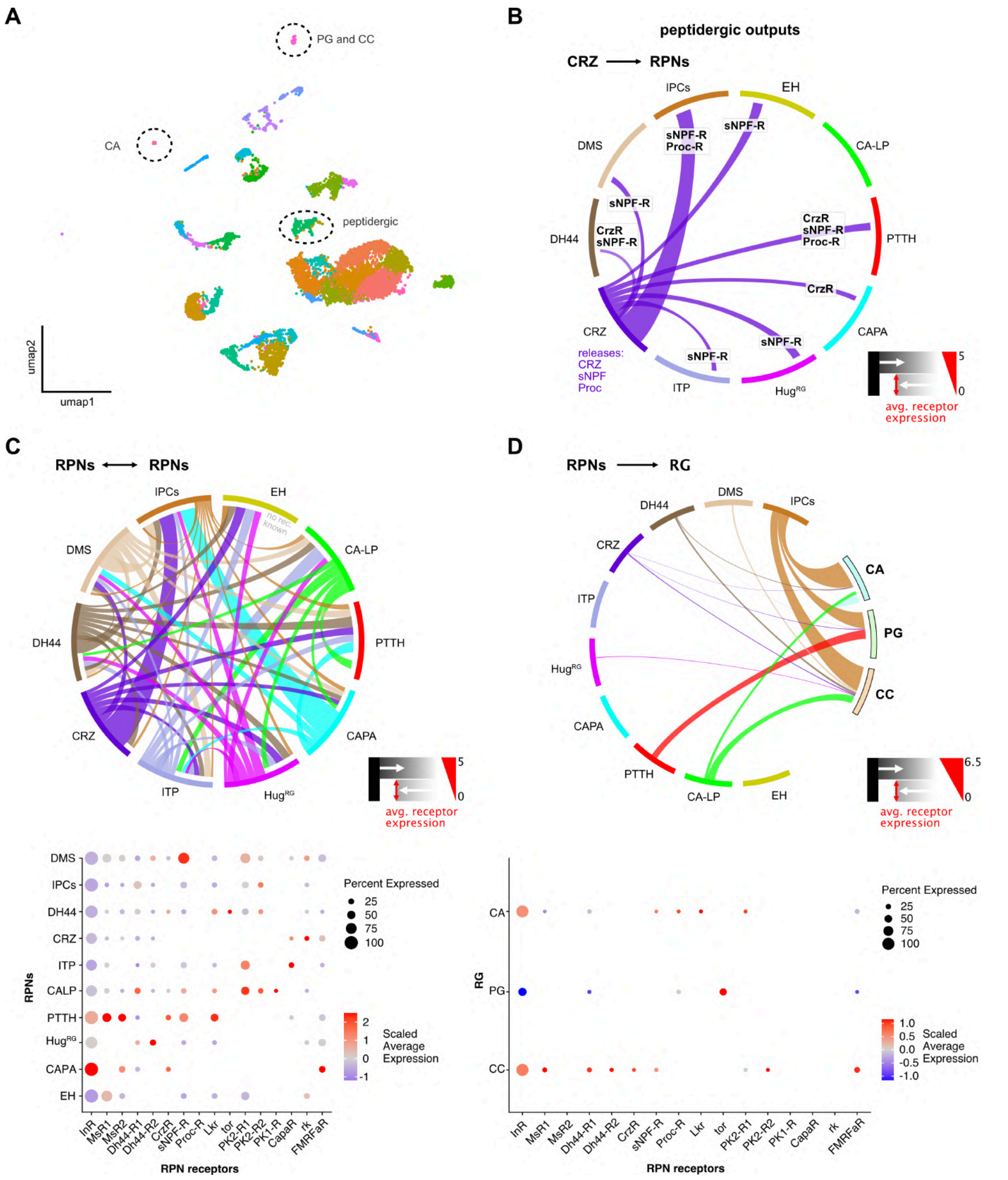


**Figure 2 - figure supplement 4. Analysis of larval brain transcriptome data**. (**A**) Cluster Plot (umap) of all cell clusters derived from the larval brain transcriptome (Brunet Avalos et al., 2019). Highlighted are peptidergic and ring gland specific neurons/cells (CC, CA and PG). (**B**) graphical representation of peptide receptor interactions of CRZ neurons with other RPNs. Since CRZ neurons are known to release additionally sNPF and Proctolin, they potentially interact with RPNs having the respective receptors for these peptides. (**C**) Peptide receptor connectivity of all RPNs using the known expressed peptide and respective receptors. This illustrates dense lateral connections of RPNs. Average receptor expression is visualized by thickness of connections (0-5). Dotplot showing the scaled average peptide receptor expression (color code) and certain percentage of the RPN cell clusters (size of dots). (**D**) Peptide receptor connectivity of the RPNs to the RG areas (CC, PG and CA). InR expression is strongest in all three tissues. Dotplot showing the scaled average peptide receptor expression (color code) and certain percentage of the distinct RG areas. Analysis was performed using a modified version of the Seurat workflow published by Satija lab (Stuart et al., 2019). For details on separation parameters of RPNs please see Materials and Methods section.

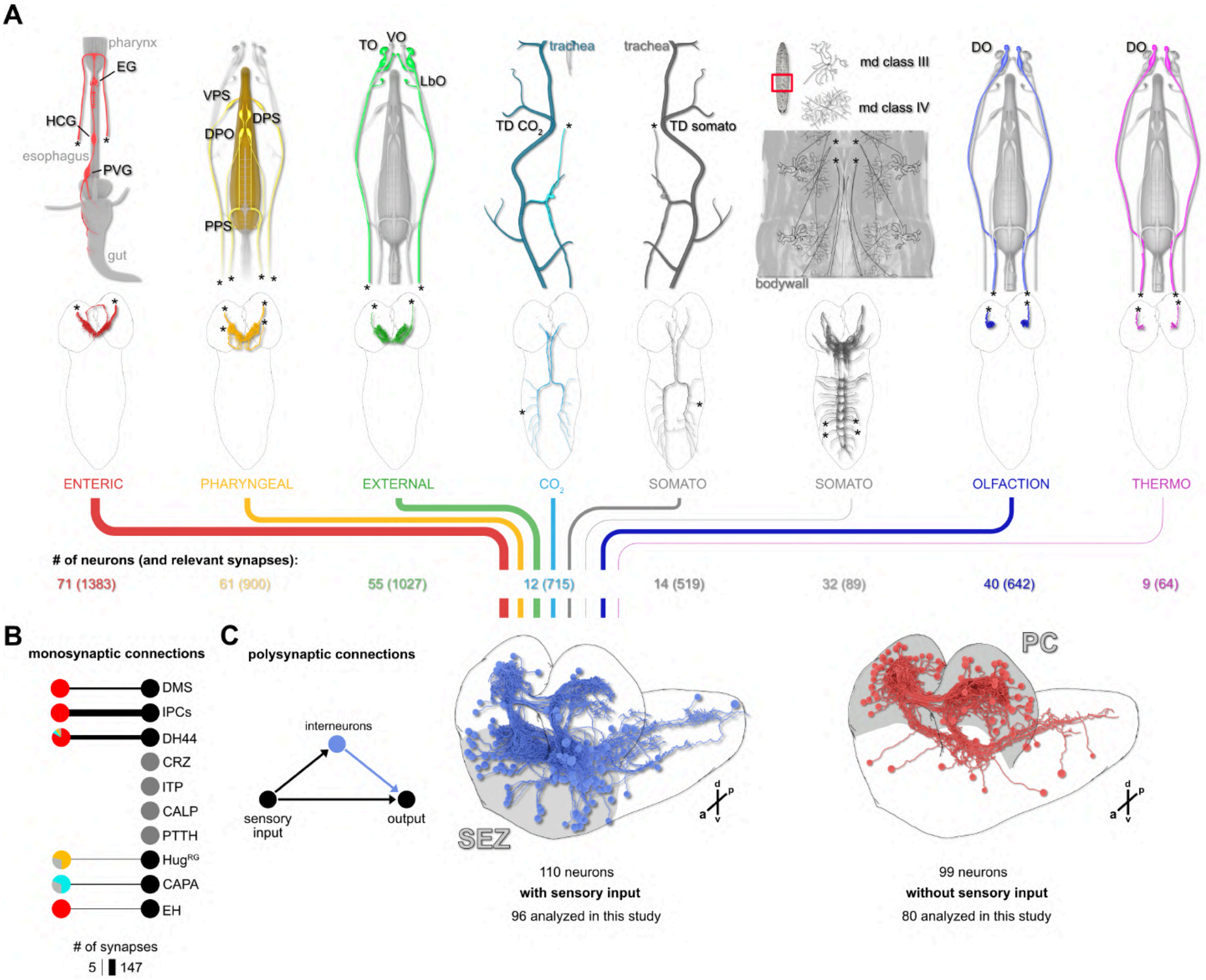


**Figure 3 - figure supplement 1. Connectivity of the distinct sensory origins to RPN upstream neurons** (**A**) *top*: schematic drawings of the dorsal view of individual sensory origins and their projections into the CNS. Strength of synaptic connections from different sensory origins is displayed as total synapse number from a given sensory origin by color coded lines (thickness represents number of synapses). Main connected sensory origins (to RPN upstream neurons) derive from enteric, pharyngeal, external, TD $CO_2$, TD somato and olfaction (**B**) all monosynaptic connections to RPNs. Note that neurons of the pars lateralis (CRZ, ITP, CA-LP and PTTH) have no monosynaptic connections to the sensory system. (**C**) Presynaptic neurons of all RPNs (threshold = 3 synapses) were divided based on connectivity to sensory neurons (min 2 synapses, blue) or having no or minor connectivity to sensory neurons (<2 synapses) (red). Analyzed interneurons: Interneurons having 3 synapses to RPNs and their hemilateral partners connected to each RPN with at least one synapse. If a hemilateral partner of an interneuron had no synaptic connection to a given RPN group the interneuron was excluded from analysis.

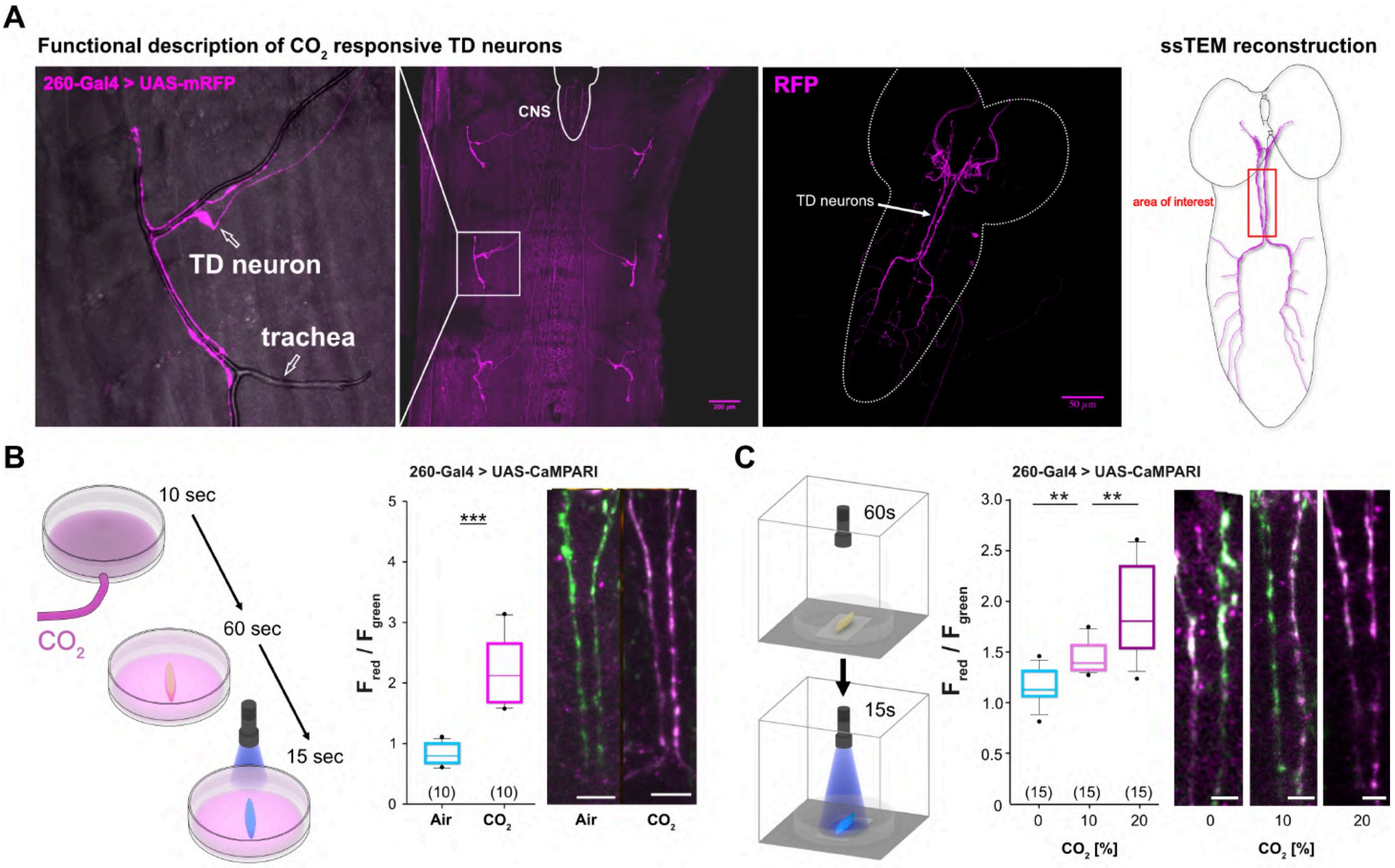


**Figure 3 - figure supplement 2. A subset of TD sensory neurons responds to $CO_2$.** (**A**) Expression of TD neurons in 260-Gal4 line crossed with UAS-mRFP. TD neurons wrap around peripheral trachea and project to the VNC and SEZ. *right*: Dorsal view of TD neurons projecting to the SEZ in the L1 ssTEM volume. Area of interest was used in 260-Gal4 line to measure $F_{red}/F_{green}$ ratio in experiments. (**B**) Schematic of experimental setup. $CO_2$ was blown for 10sec into petri dish, larva was placed in petri dish for 1 min and subsequently illuminated with 405 nm light for 15 sec. $F_{red}/F_{green}$ ratios shown for TD neuron dendrites projecting to the SEZ in 260-Gal4 line crossed with UAS-CaMPARI-1. Strong photoconversion was observed, when larvae encountered $CO_2$ enriched petri dishes. (**C**) Schematic setup for dose dependent $CO_2$ stimulation. Similar to B, but $CO_2$incubator was set to concentration of 0, 10 or 20% $CO_2$. Photoconversion of red to green fluorescence could be observed dose dependently with rising concentrations of $CO_2$ in TD neurons.

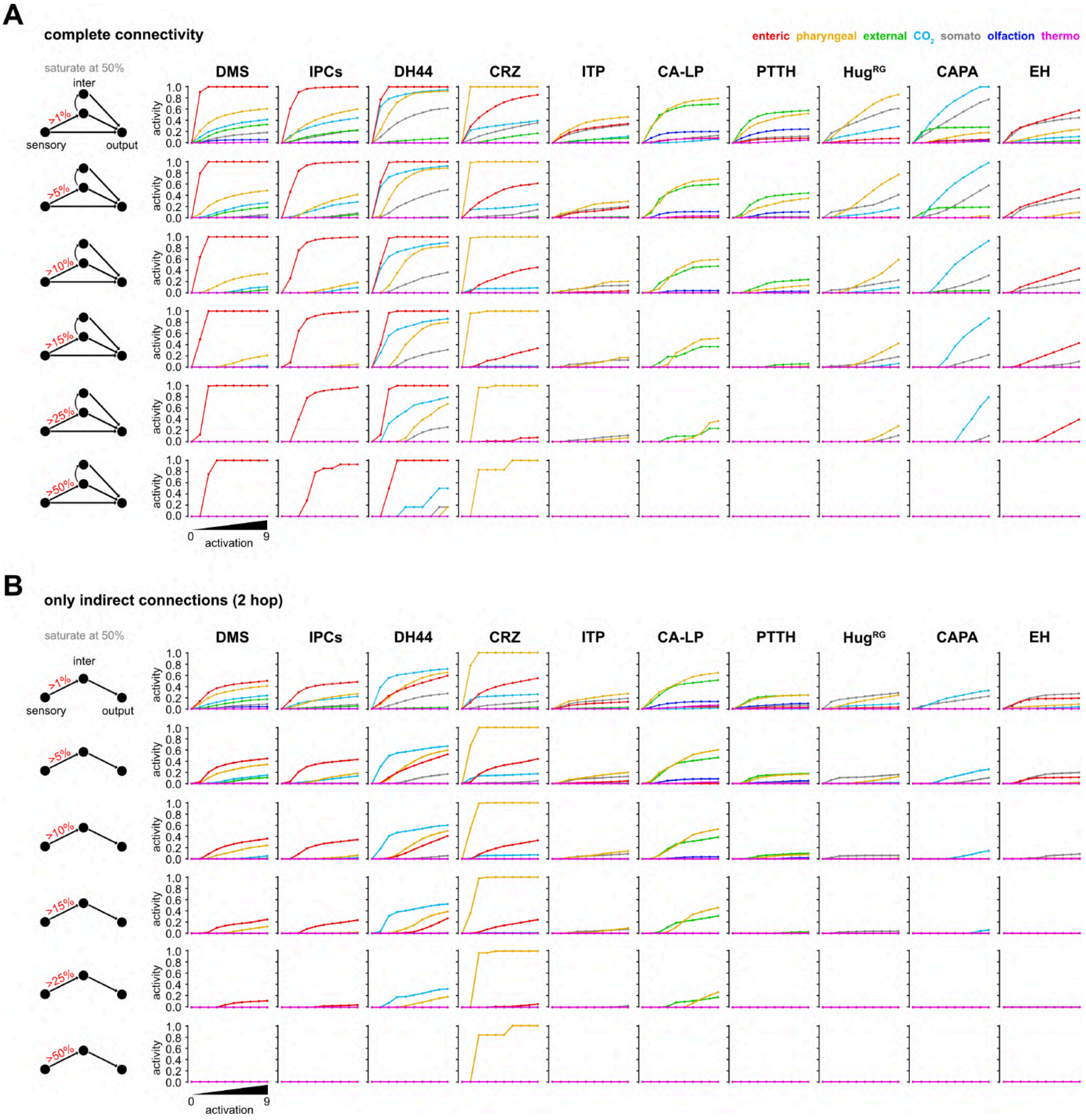


**Figure 3 - figure supplement 3.1 Parameter changes in the FFN. (A)** activity modulation of RPNs using the complete connectivity to the sensory system (monosynaptic = direct, 2-hop polysynaptic = indirect and 3-hop polysynaptic = indirect. Only interneurons were used, which have at least 3 synapses to each of the RPNs. **(B)** activity modulation of RPNs using only 2-hop indirect connections.

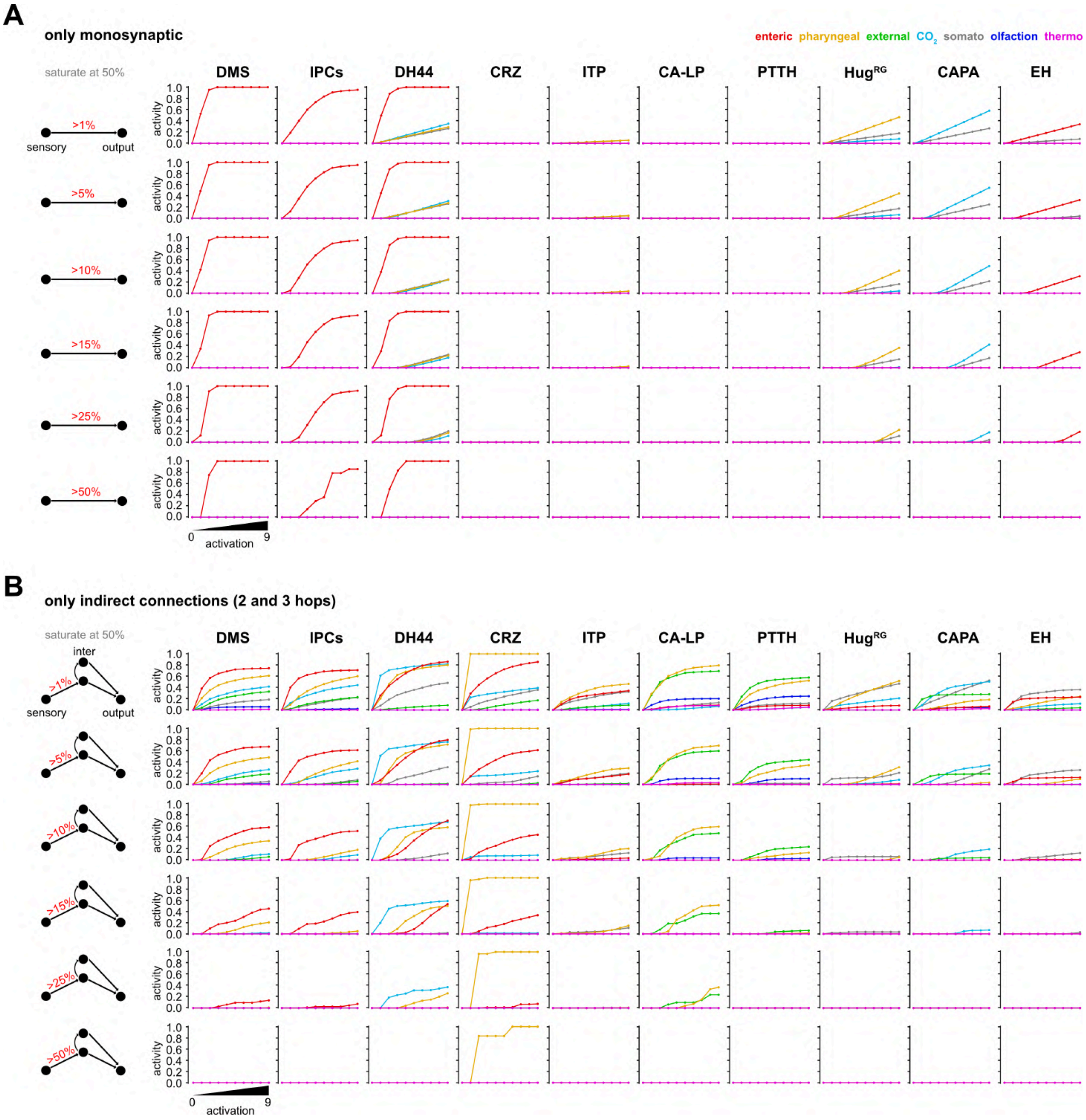


**Figure 3 - figure supplement 3.2. Parameter changes in the FFN. (A)** activity modulation of RPN using only monosynaptic connections to the sensory origins. **(B)** activity modulation of RPNs using 2- and 3-hop polysynaptic connections.

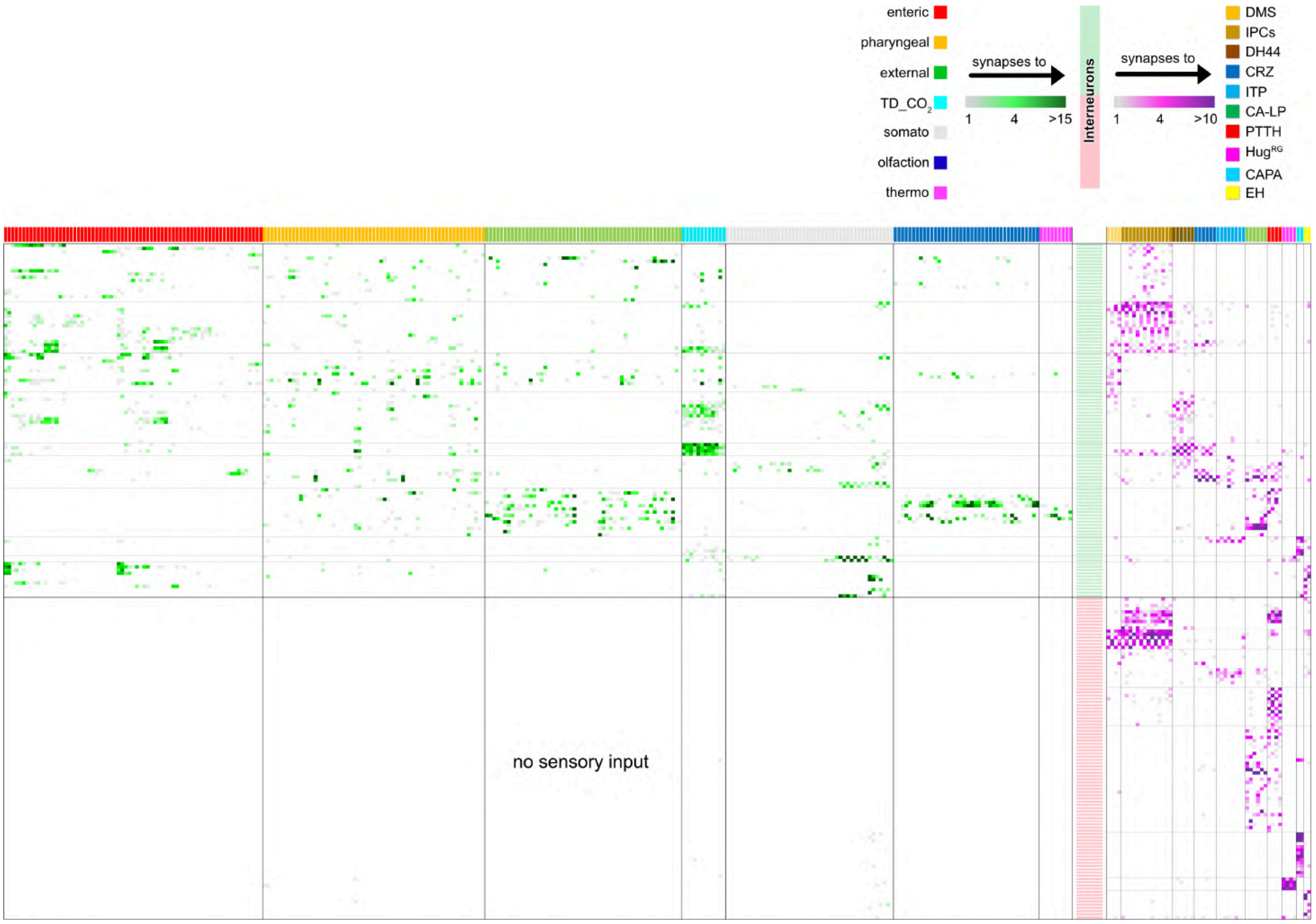


**Figure 3 - figure supplement 4. Adjacency matrix of all neurons used in this study.** Adjacency matrix showing sensory to interneuron (no synaptic threshold) and interneuron (selected with a synaptic threshold of at least 3 synapses to any of the RPNs) to RPNs. All synapses between shown neuron types were included. *top*: color coded sensory origins synapse onto upstream neurons (interneurons) of the RPNs (green heatmap). Upstream neurons synapse onto RPNs (magenta heatmap), clustered by peptide identity. Matrix: Sensory origins are displayed by single cells color coded as above scheme. Interneurons were divided into receiving and not receiving synaptic input from sensory neurons. Matrix was formatted using the connectivity of interneurons to specific peptidergic RPN clusters.

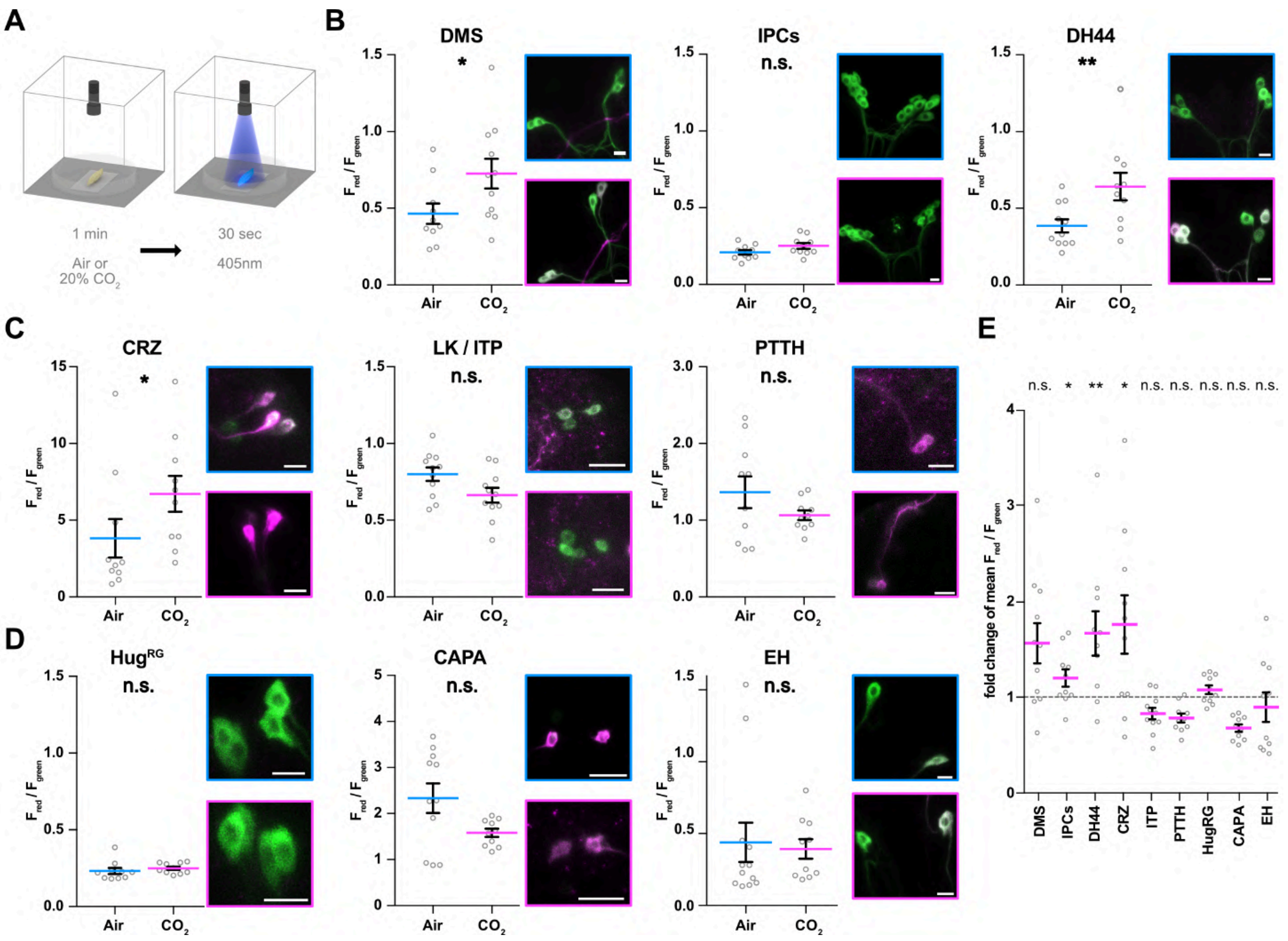


**Figure 4 - figure supplement 1.** Impact of $CO_2$ stimulation on RPNs. **(A)** Setup for $CO_2$ stimulation of intact larvae. $CO_2$ incubator with either 0 or 20% $CO_2$ was used to place larvae on a petri dish beneath 405nm photoconversion light. After 1 min of $CO_2$ exposure UV light was activated for 30 seconds. **(B-D)** Different peptide Gal4-lines expressing UAS-CaMPARI-2 in RPN clusters. Note that certain peptidergic clusters show baseline activity (CRZ, PTTH, CAPA) and therefore different scaling for the y-axis was used, which represents the red to green fluorescence ratio. Significant activity changes could be observed for DMS, DH44 and CRZ neurons upon $CO_2$ stimulation (magenta bars) compared to Air (blue bars). Images besides graphs show representative maximum projections of imaged cells (blue border = Air, magenta border = 20% $CO_2$). **(E)** Graph representation of fold change of the red to green fluorescence ratio between Air and 20% $CO_2$ for each measured RPN group. Note that CA-LP neurons could not be measured due to bad fluorescence quality.

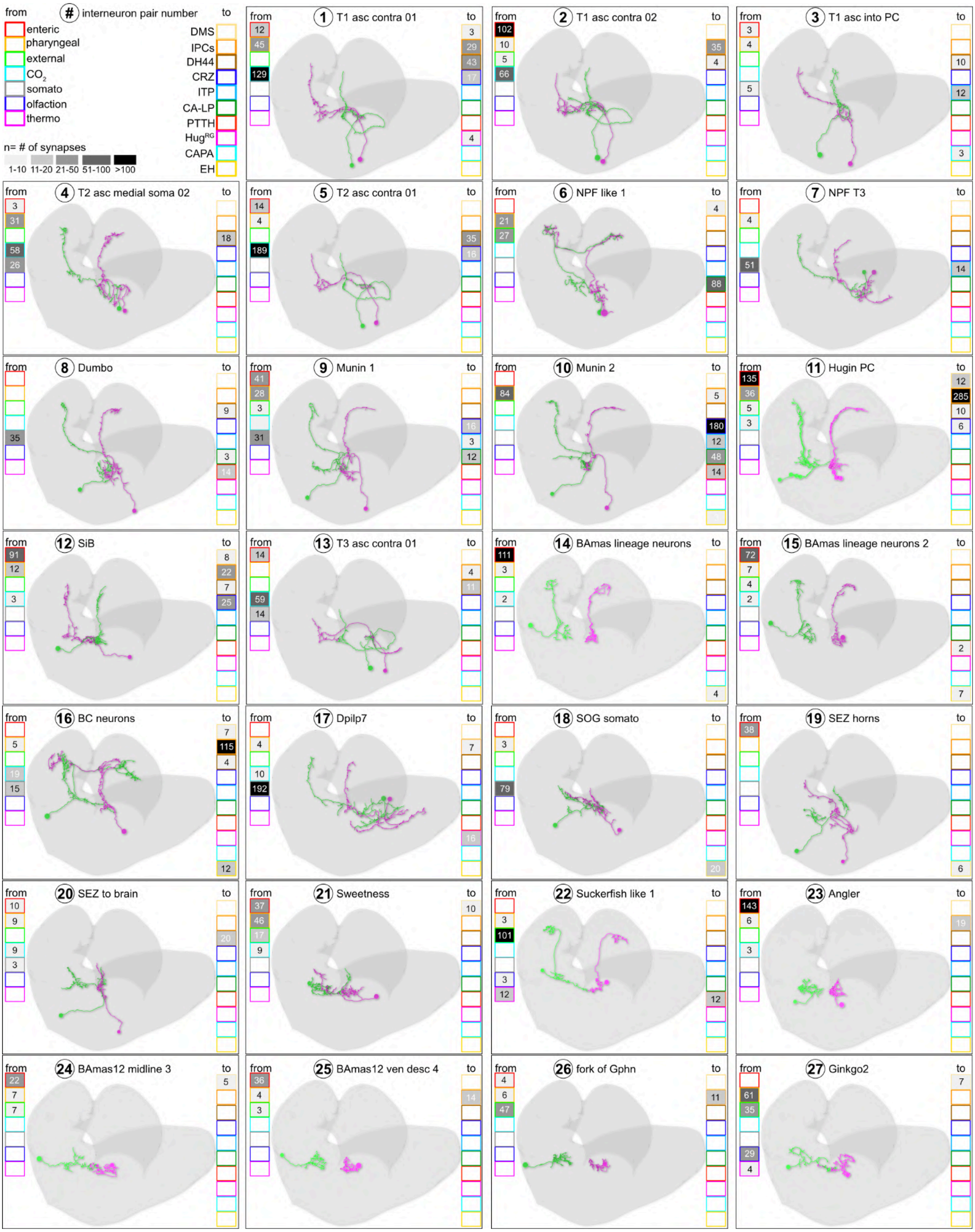


**Figure 4 - figure supplement 2.1. Neuron catalogue of interneurons receiving sensory input.** Absolute synapse numbers from sensory neurons to interneurons and from interneurons to RPN target groups are displayed in color coded boxes. Strength of synaptic contacts is heatmap coded in greyscale.

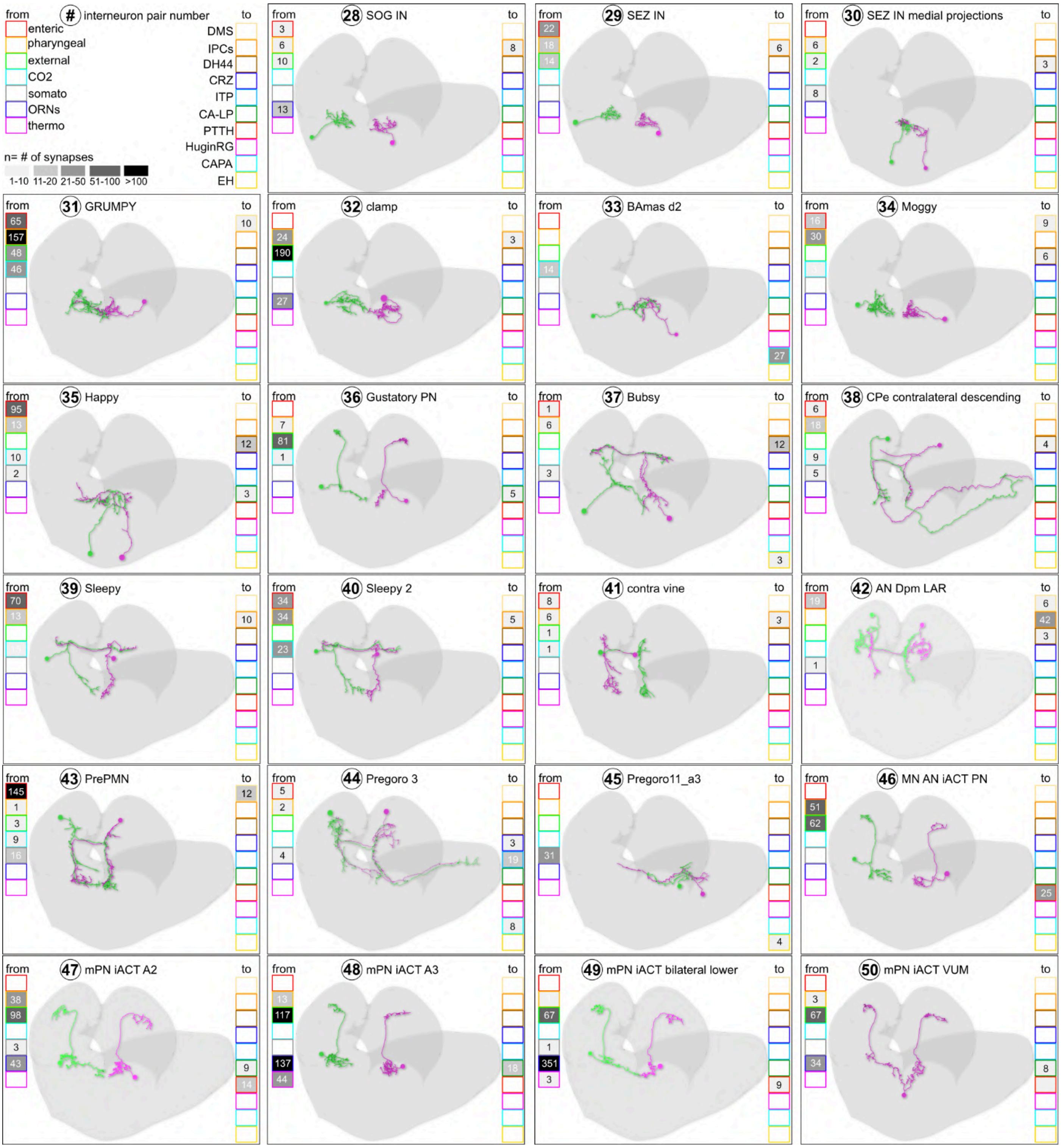

**Figure 4 - figure supplement 2.2. Neuron catalogue of interneurons receiving sensory input.** Absolute synapse numbers from sensory neurons to interneurons and from interneurons to RPN target groups are displayed in color coded boxes. Strength of synaptic contacts is heatmap coded in greyscale.

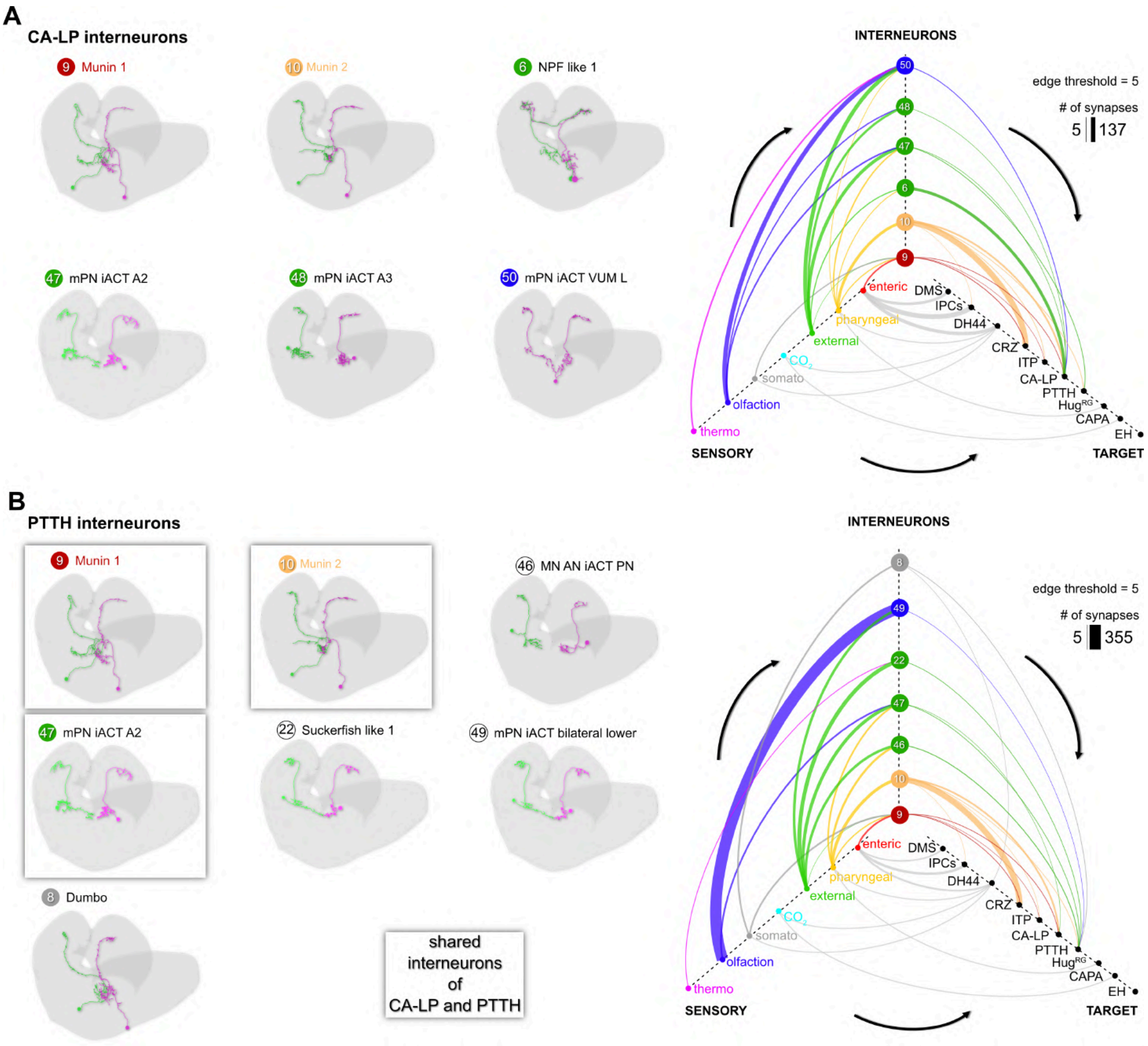


**Figure 5 - figure supplement 1. Sensory integration by CA-LP and PTTH RPNs. (A)** upstream neurons of CA-LP (synaptic threshold =3). CA-LP neurons integrate mainly pharyngeal, external and olfactory sensory signals via their upstream neurons (edge threshold =5, strongest synaptic connection has 137 synapses). **(B)** upstream neurons of PTTH. PTTH RPNs share three interneurons with CA-LP RPNs (9, 10 and 47). PTTH integrates similar sensory inputs via interneurons as CA-LP (pharyngeal, external and olfactory origins (edge threshold =5, strongest synaptic connection has 355 synapses)

.

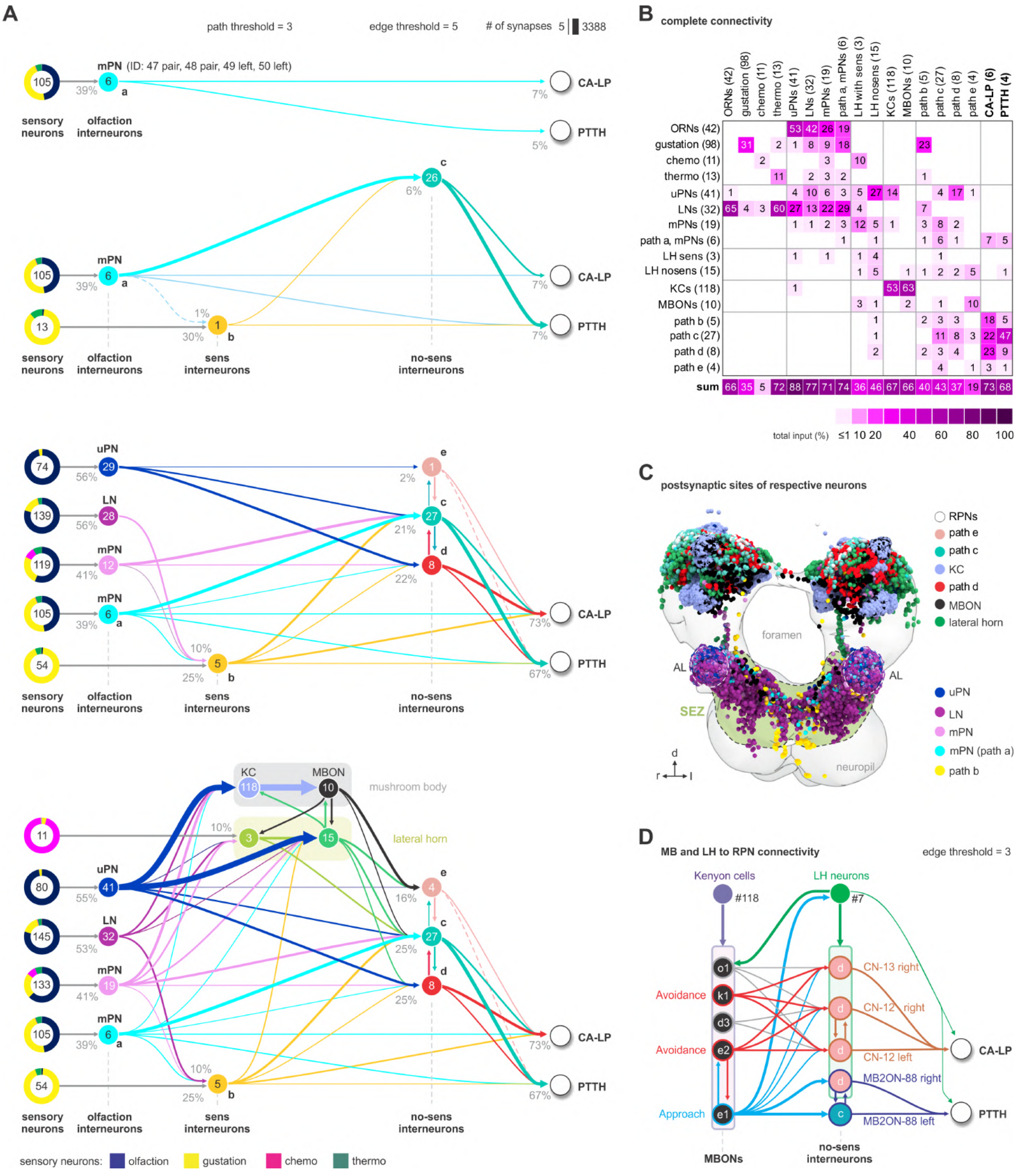


**Figure 5 - figure supplement 2. Olfactory pathways to neuroendocrine cells. (A) Panel 1.** Shortest-path (2-hops) analysis reveals six multi-glomerular projection neurons (mPNs) (Berck et al., 2016) relaying olfactory sensory information (ORNs) to CA-LP and PTTH neuroendocrine RPNs (path a). Synaptic threshold is 1 from ORNs to interneurons and minimum 3 synapses from interneurons to targets. Percentage represents the fraction of synapses from upstream neurons (arrows). Colored pie charts represent the percentage composition of total synaptic input from sensory neurons onto interneurons. Colors of pie charts correspond to the respective sensory compartments as in (Miroschnikow et al., 2018). **Panel 2.** Alternative paths (path b and c) from mPNs converging onto the same RPNs (CA-LP and PTTH) with a synaptic threshold of minimum 3. One interneuron (path b, neuron ID: 22) receives information from non-olfactory sensory neurons (sens interneurons). All 26 interneurons of path c have no connectivity to sensory neurons (no-sens interneurons). Note that path b in turn uses 4 neurons of path c as alternative routes to PRNs (syn.thresh.= 2). **Panel 3.** Adding more neurons that integrate olfactory information (olfaction interneurons): uniglomerual projection neurons (uPN), mPN and local neurons (LN)(Berck et al., 2016)) increases the number of possible and combinatorial paths (3-hops, syn. threshold = 3) to RPNs. Note that olfaction interneurons integrate information from both olfactory sensory neurons as well as from non-olfactory sensory neurons. Edges between olfaction interneurons are not shown. **Panel 4:** Superimposition of lateral horn, mushroom body kenyon cells (KC) and output neurons (MBON) onto the ORN-to-RPN circuit. Note that all sensory information-integrating neurons, mushroom body and lateral horn converge onto no-sens interneurons. **(B)** Adjacency matrix showing neurons of panel 4 in A, color-coded by percentage of inputs on targets (columns). **(C)** Frontal view of the neuropil showing the distribution of presynaptic sites of all neurons (panel 4 in A) that act as paths for olfactory information to RPNs (CA-LP-and PTTH). Each circle represents a synaptic site. Synapses are colored based on neuron class. The subesophageal zone (SEZ) and antennal lobes (AL) are marked with dotted lines. **(D)** Representation and connectivity of kenyon cells, mushroom body outputs, lateral horn neurons, convergence neurons and MB2ONs (Eichler et al., 2017; Eschbach et al., 2020) that act as paths to CA-LP and PTTH neuroendocrine RPNs (synaptic threshold = 3). Convergence neurons and MB2ON correspond to path c and e neurons (no-sens interneurons) in A. Edges of MBONs are colored based on their behavioral effect (avoidance, approach)(Eschbach et al., 2020).

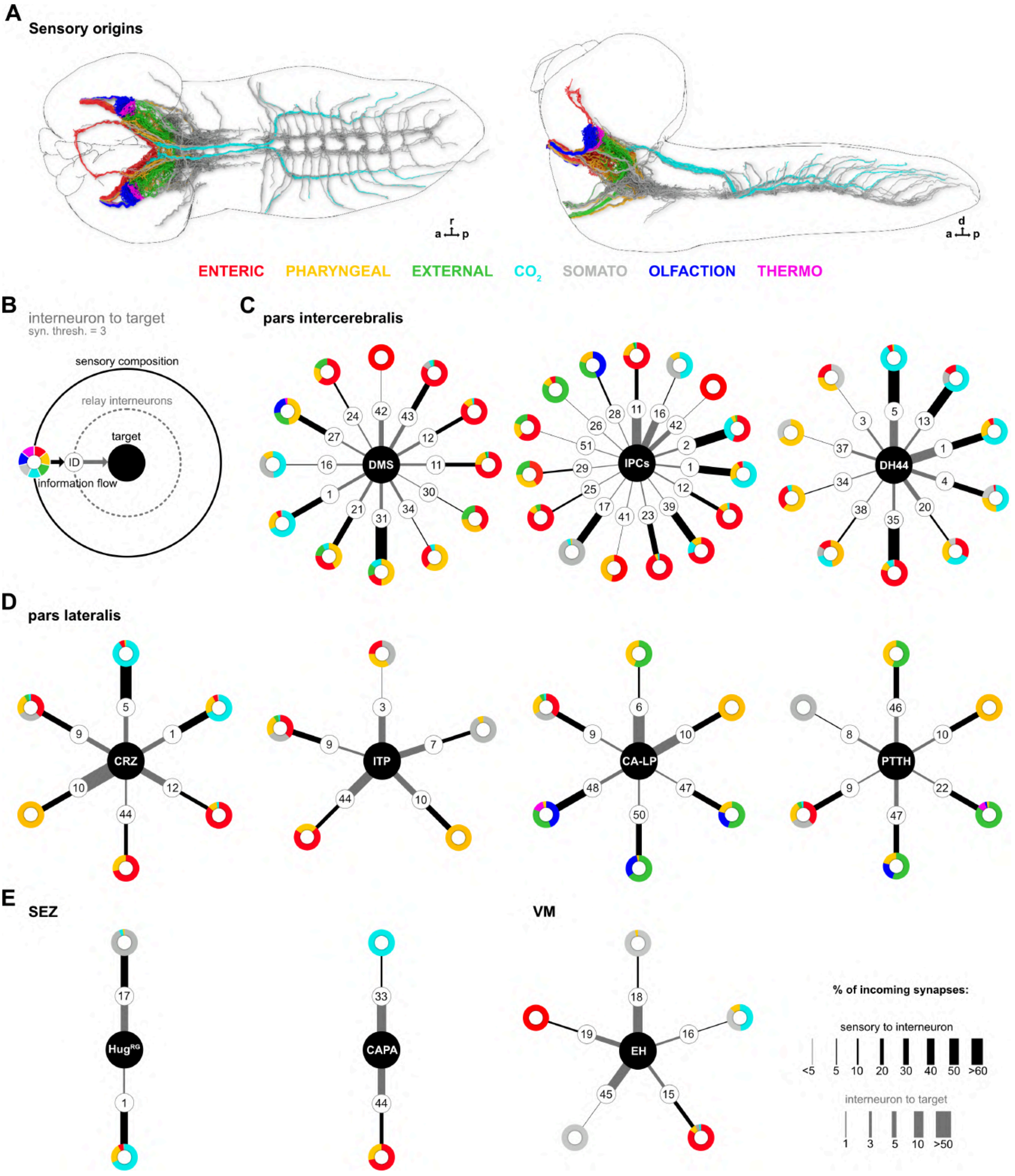


**Figure 5 - figure supplement 3. Sensory relays to RPNs of the pars intercerebralis, pars lateralis, VM and SEZ clusters. (A)** Dorsal (left) and lateral (right) view of the larval CNS and all reconstructed sensory neurons used for analysis in this study (sensory neurons were pooled from published data of earlier studies, see main text). **(B)** Schematic of graph representation. Outer "ring" represents the sensory composition of neurons targeting upstream neurons of RPNs. Line thickness to interneurons and targets represents the % of synaptic input. Striped ring represents the interneuron layer (black lined white circle). Inner ring = target neurons (RPN peptide clusters). **(C)** Sensory relay circuits of neurons within the *pars intercerebralis* region. IPCs and DMS integrate mainly information from enteric sensory areas. DH44 show most connections from TD $CO_2$ neurons via interneurons. **(D)** Sensory relay circuits within the *pars lateralis* region. Peptidergic groups show connectivity to less sensory receiving interneurons. CA-LP and PTTH are the only RPNs from PL receiving sensory information from external sensory neurons and ORNs. CRZ integrates mainly $CO_2$ and pharyngeal sensory information, while ITP neurons integrate somatosensory and pharyngeal sensory information. **(E)** Peptides of the SEZ or VM cluster show least number of sensory relay interneuron input. Hugin$^{RG}$ neurons receive sensory information from two interneuron pairs integrating mainly somatosensory and $CO_2$ sensory information; CAPA neurons integrate $CO_2$ and enteric information; EH integrates enteric and somatosensory neurons. Please note different scaling for strength of connections between sensory origin to interneurons (black lines) and interneurons to target peptide groups (grey lines).

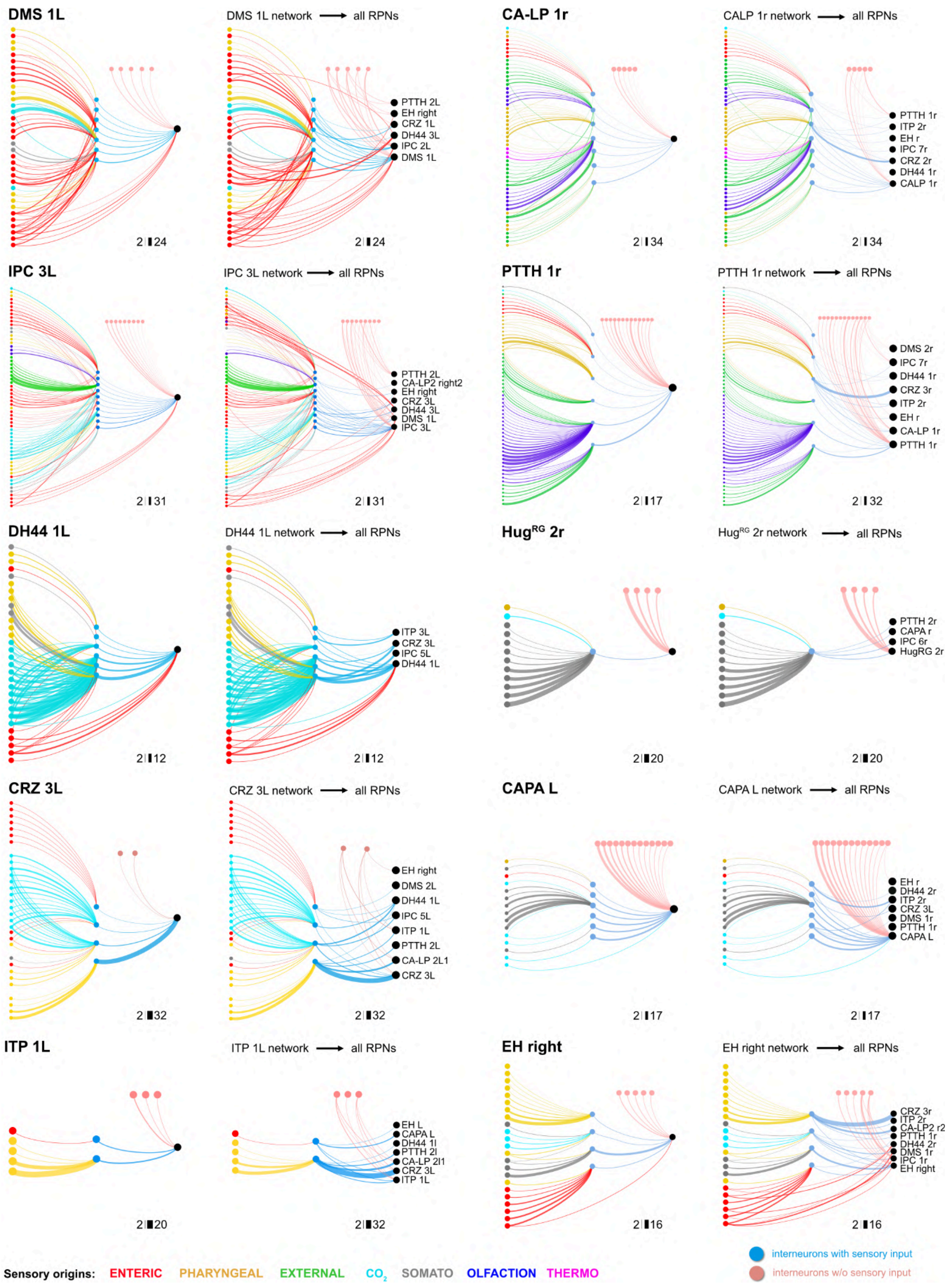


**Figure 5 - figure supplement 4**. **Diverging single cell RPN circuits.** Shown are, based on single output RPN cell, circuits from sensory origins (single cells) via interneurons (single cells) to output. Included are also interneurons (upstream of one particular RPN), which do not receive input from sensory neurons (pink interneurons). Adding additional RPNs, which are connected to the distinct interneurons, leads to diverging networks of the RPN specific single cell network to a, in strength varying, network to other RPNs.
