## Supplementary material for "Unveiling the sensory and interneuronal pathways of the neuroendocrine connectome in *Drosophila*": Figure 2 - Source Data 1

**Figure 2 - source data 1. *Drosophila* RPNs in pars intercerebralis (PI).**

| RPNs | Synonyms | Neuropeptides expressed | Neuropeptide/neurotransmitter receptors expressed | RPN Functions |
|---|---|---|---|---|
| IPCs | • mNSCs<br>• dILPs<br>• CC-PI | • Ilp1, 2, 3, 5 (Insulin-like peptide 1, 2, 3, 5) (Brogiolo et al., 2001)<br>• Dsk (Drosulfakinin) (Söderberg et al., 2012) | • Oamb (Octopamine receptor in mushroom bodies) (Crocker et al. 2010)<br>• GABA-B-R2 (metabotropic GABA-B receptor subtype 2) (Enell et al., 2010)<br>• TkR99D (Tachykinin-like receptor at 99D) (Birse et al., 2011)<br>• slo (slowpoke), Slob (Slowpoke binding protein) (Sheldon et al., 2011)<br>• CrzR (Corazonin receptor) (Kapan et al., 2012)<br>• 5-HT1A (serotonin receptor 1A) (Luo et al., 2012)<br>• AstA-R2 (Allatostatin A receptor 2) (Hentze et al., 2015)<br>• CCHa2-R (CCHamide-2 receptor) (Sano et al., 2015)<br>• PK2-R1 (Pyrokinin 2 receptor 1) (Schlegel et al., 2016)<br>• Lkr (Leucokinin receptor) (Zandawala et al., 2018)<br>• AstA-R1, -R2 (Allatostatin A receptor 1, 2), AstC-R2 (Allatostatin C receptor 2), Octβ1R (Octopamine β1 receptor) (Cocanougher et al., 2019)<br>• sNPF-R (short neuropeptide F receptor) (Oh et al., 2019) | • Diapause (Tatar and Yin, 2001)<br>• Fecundity and lifespan (Broughton et al., 2005)<br>• Sugar metabolism (Broughton et al., 2005 and 2008)<br>• Stress resistance (Broughton et al., 2005; Karpac et al., 2009; Grönke et al., 2010; Zandawala et al., 2018)<br>• Food intake regulation (Wu et al., 2005; Söderberg et al., 2012)<br>• Locomotion (Belgacem and Martin, 2006; Jones et al., 2009)<br>• Sleep (Crocker et al., 2010)<br>• Development and growth delay (Grönke et al. 2010)<br>• Sensitivity to food odors (Root et al., 2011)<br>• Starvation resistance (Kapan et al., 2012)<br>• Clock circuit (Cong et al., 2015)<br>• Food preference (Semaniuk et al., 2018) |
| DMS | • mNSCs | • Ms (Myosuppressin) (McCormick and Nichols, 1993) | • PK2-R1 (Pyrokinin 2 receptor 1) (Schlegel et al., 2016) | • Myoinhibitory on visceral, crop and gut muscles (Johnson et al., 2000; Merte and Nichols, 2002; Dickerson et al., 2012)<br>• Locomotion velocity (Kiss et al., 2013)<br>• Triger for eclosion (Ruf et al., 2017)<br>• $CO_2$ detection circuit **(this study)** |
| DH44 | • mNSCs | • Dh44 (Diuretic hormone 44) (Cabrero et al., 2002)<br>• Nplp2, 3 (Neuropeptide-like precursor 2, 3) (Cavanaugh et al., 2014)<br>• Ilp2 (Insulin-like peptide 2) (Ohhara et al., 2018) | • PK2-R1 (Pyrokinin 2 receptor 1) (Schlegel et al., 2016)<br>• Lkr (Leucokinin receptor) (Cannell et al., 2016) | • Diuretic function (Cabrero et al., 2002; Hector et al., 2009; Dus et al., 2015)<br>• Food search and feeding (Söderberg et al., 2012)<br>• Rest:activity rhythms (Cavanaugh et al., 2014)<br>• Postingestive glucose sensor (Dus et al., 2015)<br>• Sperm ejection and storage (Lee et al., 2015)<br>• Stress regulation (Cannell et al., 2016)<br>• Postingestive amino acid sensor (Yang et al., 2018)<br>• $CO_2$ detection circuit **(this study)** |

**Figure 2 - source data 1. *Drosophila* RPNs in pars lateralis (PL).**

| RPNs | Synonyms | Neuropeptides expressed | Neuropeptide/neurotransmitter receptors expressed | RPN Functions |
|---|---|---|---|---|
| CRZ | • DLP<br>• DN1<br>• CC-PL-1<br>• CN neurons | • Crz (Corazonin) (Choi et al., 2005)<br>• Proc (Proctolin) (Isaac et al., 2004)<br>• sNPF (short neuropeptide F) (Nässel et al., 2008; Kapan et al., 2012)<br>• Dsk (Drosulfakinin) (Söderberg et al., 2012) | • Dh44-R1 (Diuretic hormone 44 receptor 1), Dh31-R (Diuretic hormone 31 receptor) (Johnson et al., 2005)<br>• AstA-R2 (Allatostatin A receptor 2) (Johnson et al., 2005; Veenstra, 2009)<br>• Oamb (Octopamine receptor in mushroom bodies) (Imura et al., 2020) | • Initiation of ecdysis (Kim et al., 2004)<br>• Stress regulation (Veenstra, 2009; Kapan et al., 2012; Kubrak et al., 2016; Zhao et al., 2010)<br>• Fructose sensor (Miyamoto et al., 2012)<br>• Feeding regulation (Miyamoto et al., 2012; Hergarden et al., 2012)<br>• Ethanol tolerance (Sha et al., 2014)<br>• Egg laying (Gospocic et al., 2017)<br>• Growth regulation (via PTTH) (Imura et al., 2020)<br>• Glucose sensing/homeostasis (Oh et al., 2019)<br>• $CO_2$ detection circuit **(this study)** |
| ITP | • ipc-1<br>• ALK<br>• CC-PL-2 | • ITP (Ion transport peptide) (Dircksen et al., 2008)<br>• Lk (Leucokinin) (de Haro et al., 2010)<br>• sNPF (short neuropeptide F), Tk (Tachykinin) (Kahsai et al., 2010) | • Unknown | • Anti-diuretic, water and ion homeostasis (Kahsai et al., 2010; Gáliková et al., 2018)<br>• Food search and feeding (Gáliková et al., 2018) |
| PTTH | • PG-LP | • Ptth (Prothoracicotropic hormone) (McBrayer et al., 2007) | • CrzR (Corazonin receptor) (Imura et al., 2020) | • Regulation of ecdysone production (McBrayer et al., 2007)<br>• Promotes light avoidance at end of larval stage (Yamanaka et al., 2013)<br>• Circadian rhythmicity of eclosion (Selcho et al., 2017)<br>• Metamorphosis onset, reproductive capacity (Shimell et al., 2018)<br>• Growth (Colombani et al., 2012) |
| CA-LP | | • FMRFa (FMRFamide) (Hartenstein, 2006)<br>• Burs (Bursicon) **(this study)** | • Unknown | • Unknown |

**Figure 2 - source data 1. *Drosophila* RPNs in subesophageal zone (SEZ) or protocerebrum.**

| RPNs | Synonyms | Neuropeptides expressed | Neuropeptide/neurotransmitter receptors expressed | RPN Functions |
|---|---|---|---|---|
| HugRG | • CC-MS1 | • Hug (Hugin) (Melcher and Pankratz, 2005; Schlegel et al., 2016) | • Unknown | • Unknown function for this subpopulation of Hugin cells |
| CAPA | • CC-MS2 | • Capa (Capability) (Kean et al., 2002; Wegener et al., 2006) | • Unknown | • Unknown |
| EH | • VM neurons | • Eh (Eclosion hormone) (Hodoroyski et al., 1993) | • ETHR (Kim et al., 2006) | • Onset of ecdysis behavior (Truman, 1992; Baker et al., 1999)<br>• Coordination of eclosion (McNabb et al., 1997)<br>• Tracheal filling (Baker et al., 1999)<br>• Pre-ecdysis behavior (Krüger et al., 2015) |

Much of the information presented in Figure 2 - source data 1 is comprehensively shown and summarized in Siegmund and Korge (2001), Nässel et al. (2008), Nässel and Winther (2010), Nässel et al. (2013) and Nässel and Zandawala (2019).
